# Rice NH2 Functions as a Positive Regulator of Salicylic Acid–Mediated Defense Responses Against Sheath Blight and Bacterial Blight

**DOI:** 10.1101/2025.11.17.688593

**Authors:** Vignesh Ponnurangan, Shanthinie Ashokkumar, Krish K. Kumar, Kokiladevi Eswaran, Arul Loganathan, Sudhakar Duraialagaraja, Gopalakrishnan Chellappan, Paranidharan Vaikuntavasan, Djanaguiraman Maduraimuthu, Varanavasiappan Shanmugam

## Abstract

The rice genome encodes five non-expressors of pathogenesis-related (NPR) homologs, with OsNPR1/NH1 and OsNPR3/NH3 emerging as pivotal players in salicylic acid (SA)-mediated defense responses. Investigating the functional implications of the remaining NPR/NH genes is critical for the development of disease-resistant rice cultivars. This study explores the role of OsNH2 in rice defense against sheath blight (ShB) using CRISPR/Cas9-edited mutants of the susceptible cultivar ASD16 and the moderately resistant CO51. *OsNH2* knockout mutants showed increased susceptibility to ShB, as evidenced by dense mycelial growth, wider hyphae, and elevated superoxide radical content. Two in-frame deletion mutants lacking 15–17 amino acids in the BTB/POZ domain also showed higher susceptibility, highlighting the importance of an intact OsNH2 protein for resistance. qRT-PCR analysis revealed significant downregulation of *OsNH1*, *OsNH3*, key transcription factors (*WRKY4*, *WRKY45*, *WRKY80*, *TGA2* and *TGA3*), pathogenesis-related (PR) genes (*PR1*, *PR3* and *PR5*), and SA biosynthesis genes (*PAL* and *ICS1*) in the mutants. Additionally, *OsNH2* mutants in both cultivars exhibited reduced endogenous SA levels upon *Rhizoctonia solani* infection. Exogenous SA treatment partially restored resistance and upregulated *OsNH1/3* expression in mutants, though not to wild-type levels. These results suggest that OsNH2 is essential for maintaining SA-mediated defense signaling and optimal expression of NPR1 homologs. Moreover, *OsNH2* mutants also showed increased susceptibility to bacterial leaf blight (BLB). Collectively, this research highlights the critical role of OsNH2 in coordinating with OsNH1 and OsNH3 in SA-mediated defense against ShB and BLB in rice.

**Highlights:** CRISPR/Cas9-edited *OsNH2* knockout mutants, along with in-frame deletion mutants lacking 15–17 amino acids in the BTB/POZ domain, exhibited increased susceptibility to sheath blight disease in rice.

OsNH2 disruption led to reduced endogenous salicylic acid (SA) levels and significant downregulation of

*OsNH1*, *OsNH3*, key WRKY and TGA transcription factors, and pathogenesis-related (PR) genes.

Exogenous SA treatment partially restored resistance and upregulated *OsNH1*/*3* expression in mutants, though not to wild-type levels—highlighting OsNH2’s essential role in sustaining SA-mediated defense signaling.

*OsNH2* mutants also showed increased susceptibility to bacterial leaf blight (BLB), emphasizing its coordination with *OsNH1* and *OsNH3* in defense against multiple rice pathogens.

**Graphical abstract:** 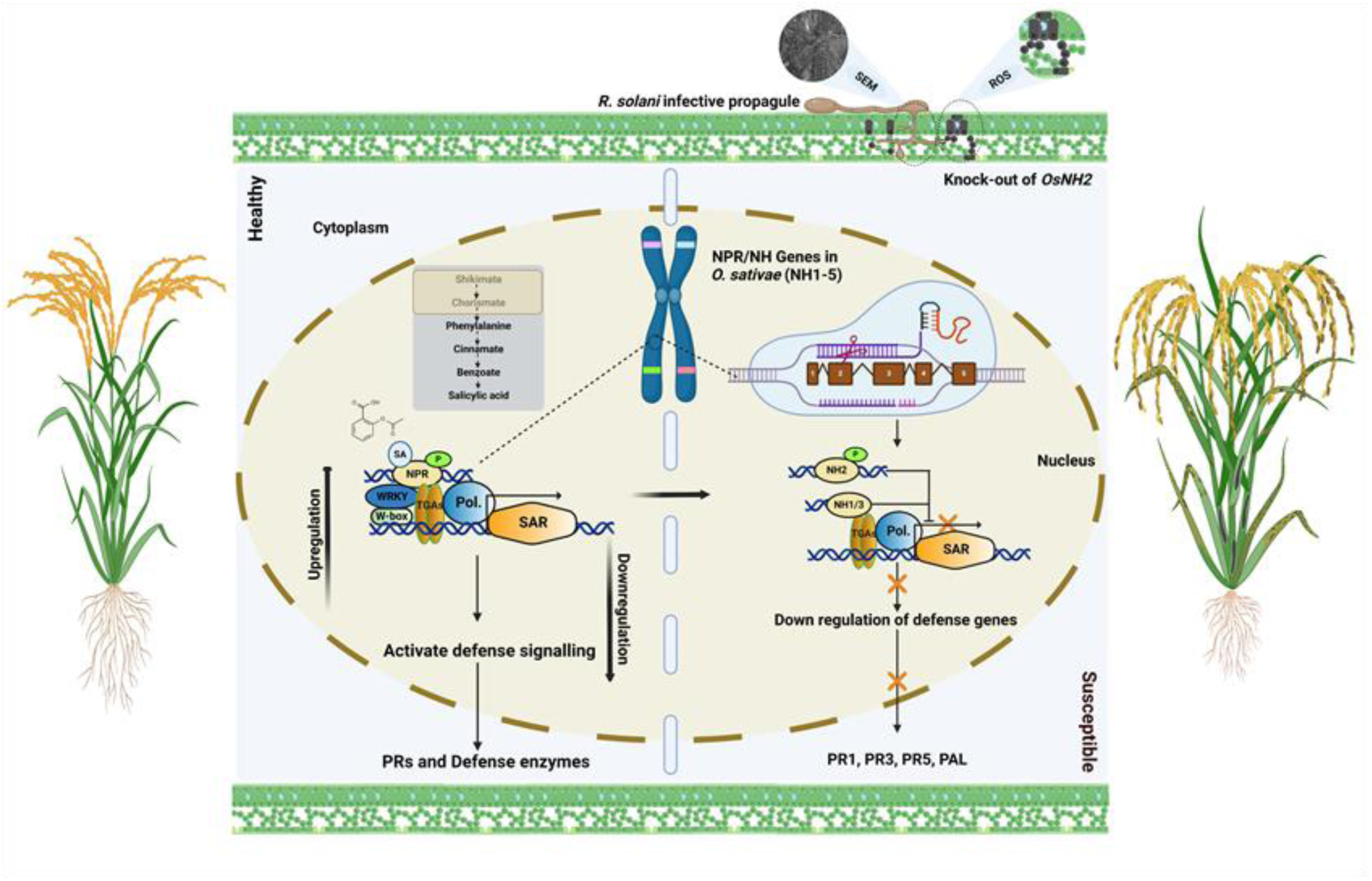

## Introduction

Rice production depends on several biotic and abiotic factors, which can significantly affect crop yield and grain quality. Among these, biotic factors have a profound effect on rice yield and quality, posing a major challenge to sustainable rice cultivation (Senapati et al., 2022). Several fungal and bacterial diseases pose significant threats to rice cultivation. One of the major fungal diseases affecting rice is sheath blight (ShB) is caused by *Rhizoctonia solani*, a soilborne necrotrophic fungus that produces sclerotia of different sizes, which can remain dormant for many years (Molla et al., 2020). Depending on the severity of the infection, ShB can cause a yield reduction ranging from 20 to 50% (Margani and Widadi, 2018; Ponnurangan et al., 2025). Previous studies suggest that the breakdown of ShB disease resistance is attributable to the pathogen’s rapid evolution, leading to severe disease outbreaks (Zheng et al., 2013). It has been reported that generating host plant resistance against ShB can result in an 8% yield gain in rice (Willocquet et al., 2004). However, one of the biggest challenges in rice breeding is identifying new genes that resist biotic stresses (Jesudoss et al., 2024).

In plants, nonexpressor of pathogenesis-related (NPR) genes play a crucial role in immunity, particularly in systemic acquired resistance (SAR). In *Arabidopsis*, NPR1, also known as *NIM1* or *SAI1*, has been identified as a key positive regulator of the salicylic acid (SA)-dependent signaling pathway (Cao et al., 1994). NPR1 plays a significant role in regulating broad-spectrum resistance in plants, as SA binds to NPR1, thereby enhancing the activation of defense gene transcriptional activity, which leads to increased plant disease resistance (Backer et al., 2019). The NPR1 protein comprises an N-terminal BTB/POZ (broad-complex, tram-track, and bric-a-brac/pox virus and zinc finger) domain, a central ankyrin repeat domain, and an NPR1/NIM1-like defense domain, all of which are crucial for protein interactions that regulate plant defense responses (Cao et al., 1997; Rochon et al., 2006). NPR1 and its homologs have been identified in numerous plants, including rice, and their role in disease resistance has been investigated. In *Arabidopsis*, paralogs of *AtNPR1*, namely NPR3 and NPR4, share very similar domain structures (Cao et al., 1998; Zhang et al., 2006).

In contrast to *AtNPR1*, *AtNPR3* and *AtNPR4* act as transcriptional co-repressors in plant defense mechanisms, playing a negative regulatory role in SA-mediated disease signaling events. Studies have shown that targeted mutations in NPR3/4 enhance immune responses to bacterial and fungal infections in *Arabidopsis* and cacao, thereby confirming the negative regulatory role of these proteins in plant disease resistance (Fister et al., 2018; Ding et al., 2018). The SA receptors NPR1 and NPR3/NPR4 play opposing roles in transcriptional regulation during plant defense mechanisms. Arabidopsis NPR3/NPR4 suppresses defense-related gene expression by interacting with TGA2/TGA5 at low SA levels, but releases the transcriptional repression at increased SA levels, while the regulatory functions of both NPR1 and NPR3/NPR4 depend on interactions with TGA transcription factors (Ding et al., 2018). NPR1 also interacts with WRKY transcription factors in regulating SA-mediated plant defense response (Chen et al., 2020; Yu et al., 2001).

In the rice genome, six genes have been identified that encode five NPR1-like proteins: NH1, NH2, NH3, NH4, and NH5 (NPR1 homologs 1–5). The NH genes are categorized into three Clades: *OsNH1* belongs to Clade 1, *OsNH2* and *OsNH3* to Clade 2, and *OsNH4* and *OsNH5* to Clade 3 (Yuan et al., 2007; Bai et al., 2011; Chern et al., 2014; Moon et al., 2018). OsNH1/OsNPR1 is the closest relative to AtNPR1, exhibiting 58% amino acid identity. The amino acid sequences of NH2 and NH3 share 54% identity, whereas those of NH4 and NH5 show 62% identity, confirming that rice NPR1-like proteins are closely related. Sequence identity analysis of NH2/NH3 and NH4/NH5 reveals a close evolutionary relationship between these proteins, suggesting potential similarities in their roles (Bai et al., 2011). In rice, the overexpression of *AtNPR1/OsNH1* leads to increased resistance against pathogens such as *Xanthomonas oryzae* pv*. oryzae* (*Xoo*) and *Magnaporthe grisea* (Chern et al., 2005; Yuan et al., 2007). Transgenic rice lines overexpressing *OsNPR2* and *OsNPR3* under the constitutive CaMV 35S promoter did not exhibit enhanced resistance to *Xoo*. In contrast, introducing an additional copy of *OsNH3* driven by its native promoter resulted in significantly enhanced resistance against *Xoo* (Bai et al., 2011). The mechanisms by which NH1 and NH3 confer immune responses are well understood, whereas the roles of other genes in the same group—NH2, NH4, and NH5— remain unclear (Jun et al., 2010; Chern et al., 2014). However, a recent study demonstrated that overexpression of a novel NH5 gene, *OsNH5N16*, conferred enhanced resistance to BLB and Bakanae disease by upregulating PR genes associated with SAR (Son et al., 2021). Among the NPR gene family members, the role of *OsNH2* in rice disease resistance has not been clearly established.

In the present study, the CRISPR/Cas9 technology was used to knock out *OsNH2* and subsequently understand its function in ShB resistance. To investigate the role of *OsNH2*, targeted mutations were carried out in the highly susceptible cultivar ASD16 and the moderately resistant cultivar CO51. In addition, pathogenicity assays with the bacterial leaf blight (BLB) pathogen were carried out further to explore the role of *OsNH2* in disease resistance mechanisms. Disease screening against ShB and BLB revealed that mutants from both rice cultivars showed higher susceptibility than wild-type (WT) plants. This enhanced susceptibility of *OsNH2* mutants to both fungal and bacterial pathogens reveals the critical role of *OsNH2* in rice defense responses, highlighting *OsNH2* as a key component in conferring resistance against multiple pathogens.

## Materials and Methods

### Construct preparation and rice genetic transformation

The *OsNH2* gene is located in the 1st chromosome of rice, and its sequence information (Locus ID BGIOSGA004522), approximately 3.13 kb in length with five exons, was retrieved from the Ensembl Plants database (http://plants.ensembl.org). The single-guide RNA (sgRNA) was designed using the web-based CRISPR-P v2.0 tool (http://cbi.hzau.edu.cn/crispr/). To facilitate cloning at the *Bsa*I restriction site of pRGEB32 (Addgene plasmid # 63142) (Xie et al., 2015), the following adaptors were added to the oligos: top strand 5’-GGCA-3’ and bottom strand 5’-AAAC-3’. The recombinant plasmid was cloned into *Escherichia coli* and subsequently mobilized into *Agrobacterium tumefaciens* (LBA4404) *via* triparental mating. A positive transconjugant colony was used for the cocultivation of immature embryos. Putative mutants were generated in cultivars of ASD16 and CO51 genetic backgrounds using *Agrobacterium-*mediated genetic transformation of immature embryos, following the protocol of Hiei and Komari (2008). The putative mutants developed were hardened and maintained in a transgenic greenhouse.

#### Molecular characterization of *OsNH2* mutants

Plant genomic DNA was isolated from putative *OsNH2* mutants and WT cultivars of ASD16 and CO51 using the CTAB method (Porebski et al., 1997). Marker genes such as *hpt* and *Cas9* were confirmed using PCR. Target-specific primers were used to amplify the sgRNA region, which was then subjected to Sanger sequencing (Biokart, Bengaluru) (Table S1). The sequencing results were analyzed using web tools such as DSDecodeM (Liu et al., 2015) and CRISP-ID (Dehairs et al., 2016). The *OsNH2* lines (T_0_) with mutations were selected and passed on to the T_1_ generation to identify homozygous mutations. Seeds were harvested from homozygous *OsNH2* mutants and used for bioassay studies. The nucleotide sequences of homozygous *OsNH2* mutants were translated into protein sequences using the online bioinformatics tool ExPASy (https://www.expasy.org) and aligned with the WT amino acid sequence using Clustal Omega (https://www.ebi.ac.uk/Tools/msa/clustalo/).

The putative off-target loci specific to the selected *OsNH2*-sgRNA sequence were predicted using online tools, namely CRISPR-P v2.0 (Liu et al., 2017) and CRISPR-GE (Xie et al., 2017). Four off-target sites were identified for each sgRNA, selected based on factors such as high sensitivity, a maximum of four mismatches, and location predominance in coding regions (Table S2). The genomic regions encompassing these predicted off-target sites were amplified by PCR using site-specific primers (Table S1), followed by Sanger sequencing of both the *OsNH2* mutants and the WT (Xu et al., 2015; Shanthinie et al., 2024). Also, the off-target sites in other members of the NH gene family were sequenced to assess any unintended edits (Chern et al., 2014). The sequences were aligned with the WT sequence to identify any off-target edits in the homozygous *OsNH2* mutants.

#### Growth conditions for pathogens and plant materials

*R. solani* AG1-1A was grown on a potato dextrose agar (PDA) medium at 28°C. Mycelial discs of 3–5 mm diameter were acquired from peripheral young white mycelia of 3- to 4-day-old fungal culture using a sterile cork borer. These mycelial discs were used as the fungal inoculum in infection assays (Lin et al., 2021). For BLB inoculation studies, the *Xoo* strain was used. The culture was continuously subcultured on a nutrient agar medium and maintained at 28°C (Sree et al., 2023). Bioassay studies were carried out on CRISPR-edited *OsNH2* mutants in ASD16 and CO51 backgrounds, along with WT plants. All rice plants were grown in a greenhouse under controlled environmental conditions.

### Pathogen inoculation

#### *R. solani* inoculation

To carry out detached leaf assays, the second leaf of the main tiller was cut into 4- to 5-cm bits and placed on wet filter paper on Petri dishes. The fungal plug was excised from the PDA plate and placed on the abaxial region of the leaf. The leaves were cultured for 72 h at 25°C, and the moisture of the filter paper was maintained with sterile water (Molla et al., 2013; Gao et al., 2021). In whole-plant bioassays, a 3-day-old culture of the pathogen was inoculated onto healthy tillers, which were then covered with aluminum foil. In the control treatment, only agar plugs were used. The lesion area was measured seven days after infection (Park et al., 2008; Naeimi et al., 2020).

#### *Xoo* inoculation

A fresh *Xoo* suspension was prepared by suspending a loopful of culture in a 10 mM sterile MgCl_2_ solution. The optical density of the bacterial suspension was adjusted to 0.5 at 600 nm using a spectrophotometer (Ke et al., 2017). Homozygous mutants were inoculated with the bacterial suspension using sterilized scissors following the leaf clipping method (Kauffman et al., 1973). Sterilized scissors dipped in MgCl_2_ without *Xoo* suspension were used to cut the leaves, serving as the negative control. The plants were kept under controlled environmental conditions, and lesion length was recorded at 14 days post-inoculation (dpi) (Zhou et al., 2022; Sree et al., 2023).

#### NBT staining and scanning electron microscope (SEM) analysis

The presence of superoxide (O_2_^−^) was detected in rice leaves of mutants and WT plants, which were then infected with a 3-day-old mycelial culture of *R. solani*. Superoxide levels were visually detected using nitroblue tetrazolium (NBT) as described previously (Yang et al., 2004). Briefly, 50mM sodium phosphate buffer (pH 7.0) containing 0.2% fresh NBT solution was prepared, and 48 h after infection, the leaves were immersed in the fresh solution, covered with aluminum foil, and kept in the dark for 8 h. The leaves were then decolorized in 95% ethanol in a boiling water bath at 60°C and stored in 80% glycerol. The production of O_2_^−^ was confirmed by the appearance of blue formazan in the tissue. Superoxide staining was carried out in triplicate, which showed similar results. At least six leaves were used in each treatment.

Images of mycelial hyphae were obtained using the Quanta 250 SEM (FEI, Czech Republic). Mutant and WT leaf samples were selected after 48 h of *R. solani* inoculation and coated with gold nanoparticles using a sputter coater to enhance conductivity and improve imaging quality. Then, the samples were freeze-dried by lyophilization to maintain their structural integrity. The dried samples were then kept in the vacuum chamber of the SEM, and the imaging was conducted at a voltage of 15 kV with the sample positioned 4 mm away from the necrotic area of the leaf samples (Basu et al., 2016).

#### RNA isolation and RT-qPCR analysis

The *OsNH2* mutants and WT plants were inoculated with a 3-day-old *R. solani* culture on the sheath portion of 45-day-old rice seedlings. Samples were collected at different time intervals: 0, 24, and 72 h post-inoculation (hpi), with sheath samples from uninoculated plants serving as controls. The plants were sprayed with SA and inoculated with *R. solani* 24 h later, while the control plants were sprayed with water (Kouzai et al., 2016). The samples were quickly frozen using liquid nitrogen and stored at −80°C until further use. Total RNA was extracted using the TRIzol reagent (Sigma-Aldrich, Germany), and the concentration of RNA was determined using a NanoDrop spectrophotometer. Subsequently, complementary DNA (cDNA) was synthesized from the RNA using a RevertAid First Strand cDNA Synthesis Kit (Thermo Fisher Scientific, USA; K1622) in accordance with the manufacturer’s instructions, and the synthesized cDNA was stored at −20°C. RT-PCR was conducted (CFX Connect, BioRad) under optimized conditions using the ubiquitin gene as the reference gene. Each PCR was prepared with a total volume of 10 μl, consisting of 5 μl of Universal SYBR Green Supermix (BioRad, 1725271), 2 pmol each of forward and reverse primers, and 200 ng of cDNA samples. The PCR process involved initial denaturation at 96°C for 20 sec, followed by denaturation at 96°C for 5 sec, annealing at 60°C for 10 sec, and extension at 72°C for 15 sec with a total of 40 cycles. Melt curve analysis was carried out between 65°C and 95°C. Quantitative analysis of the results was carried out using the 2^(−ΔCt)^ method (Livak and Schmittgen, 2001; Shanmugam et al., 2015), and each reaction was run in triplicate to ensure accuracy and reliability.

#### Quantification of SA by UHPLC–MS/MS

The extraction and determination of SA were carried out following the method of Farooq et al., (2009), with slight modifications. Plant tissues were harvested at different time-course intervals as indicated, freeze-dried using a lyophilizer, and ground into a fine powder. For extraction, 100 mg of powdered tissue was mixed with 1 mL of 100% HPLC-grade methanol, vortexed for 5 min, and sonicated for 30 min at room temperature. After centrifugation at 12,000 rpm for 10 min at 4°C, the supernatant was filtered through a 0.22 μm PTFE membrane filter. SA was quantified using a Shimadzu LCMS-8045 triple quadrupole mass spectrometer coupled with a UHPLC system. Chromatographic separation was performed on a C18 column using isocratic elution with 0.1% formic acid in water and acetonitrile (60:40, v/v) at a flow rate of 0.25 mL min^-1^.

#### Evaluating the efficacy of phytohormones upon *R. solani* inoculation

Salicylic acid (Sigma-Aldrich, Cat No. 247588), jasmonic acid (JA) (Sigma-Aldrich, Cat No. 14631), and ethylene (ET) were used as phytohormones to evaluate their effects on both ASD16 and CO51 WT and *OsNH2* mutants upon *R. solani* infection. Phytohormones were dissolved in DMSO at concentrations of 0.5 mM and 1 mM, and 0.04% (v/v) Tween 20 was added to enhance foliar application. The solutions were applied as a foliar spray on 45-day-old rice seedlings, with mock-treated plants serving as the control. After 24 h of foliar spray, the mycelial discs of the *R. solani* were placed in the center of the detached leaves, which were kept moist in a Petri dish under dark conditions for 2 days. The lesion areas were then photographed and measured using ImageJ software (Kouzai et al., 2018; Wang et al., 2022).

#### Statistical analyses

Statistical analyses were conducted using R software (version 4.2.3). Differences in mean lesion length were assessed using one-way ANOVA followed by Tukey’s honestly significant difference test. Student’s t-test was used to compare significant differences in gene expression studies between controls and mutants.

## Results

### Development of *OsNH2* knockouts by CRISPR/Cas9 technology and their molecular characterization

The rice genome encodes five NPR1-like genes (NPR1 homologs 1–5), with *OsNH1* and *OsNH3* demonstrated to play roles in disease resistance (Chern et al., 2014; Yuan et al., 2007; Bai et al., 2011). This study aims to investigate the role of *OsNH2* in ShB disease resistance using the CRISPR/Cas9 technology. The rice cultivars ASD16 (highly susceptible) and CO51 (moderately resistant) were selected. Two sgRNAs targeting the 3’ end of exon 2 of the *OsNH2* gene were cloned into the pRGEB32 vector, and the constructs were designated as *OsNH2*-398 and *OsNH2*-426 (Fig. 1a). *Agrobacterium-*mediated transformation resulted in 60 and 15 independent events in ASD16 and CO51, respectively. PCR analysis confirmed the presence of *Cas9* and *hpt* genes. Sequencing of the *OsNH2* gene target region identified mutations in 39 out of 75 independent events from both cultivars, with an average mutation efficiency of 56.39% (Table S3). Due to the presence of identical mutations among these 39 events, 18 mutants were further selected, their T_1_ progeny were raised, and sequencing was performed to identify homozygous mutants (Tables S4 and S5). Among the 18 mutants, five frameshift homozygous mutants (three from ASD16 and two from CO51) and two in-frame homozygous mutants (ASD16-15/1-3 and CO51-4/2-1) were selected (Figs. 1b and c). Stable inheritance of the mutation was further confirmed in the next generation.

**Fig. 1.**
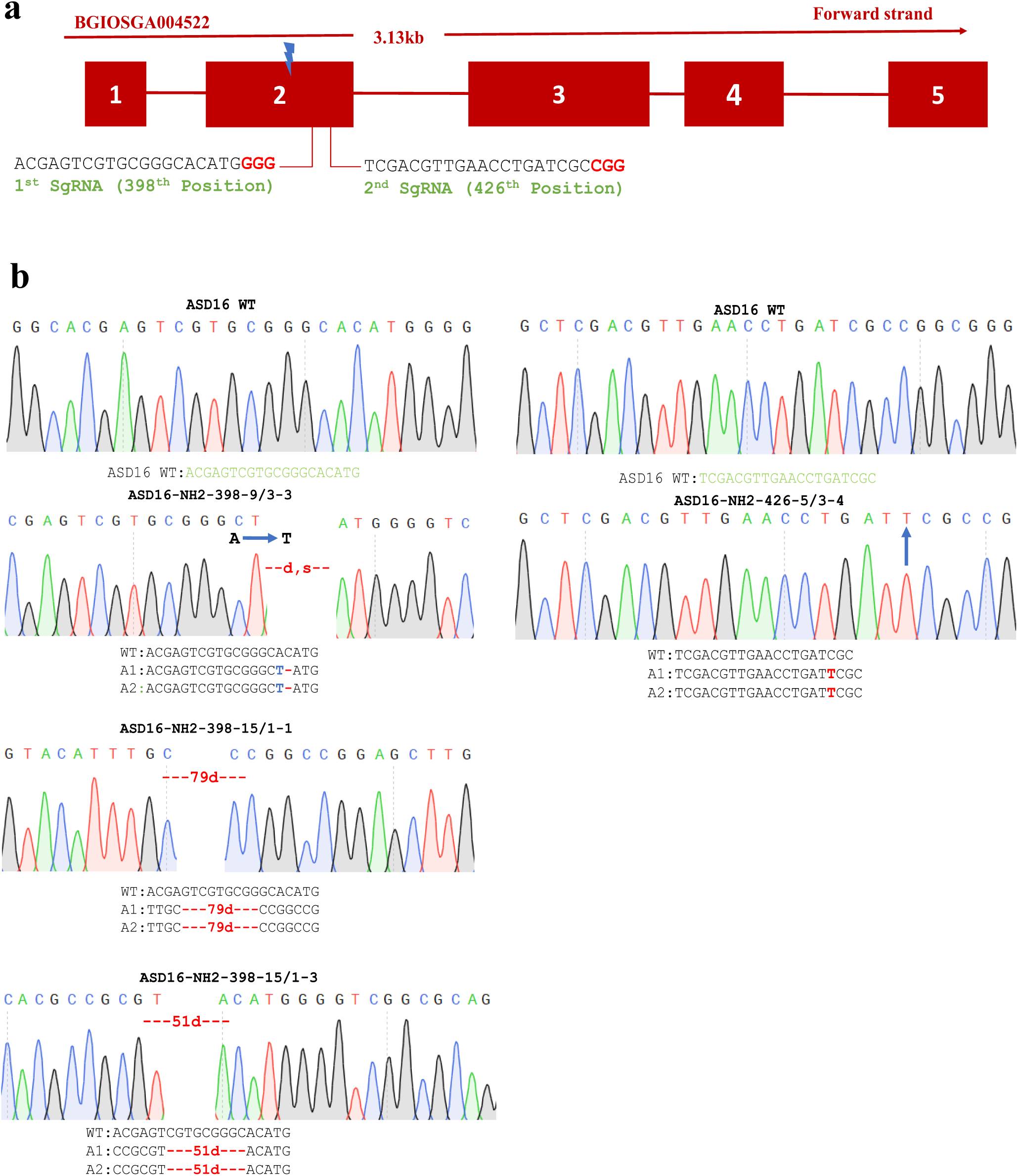
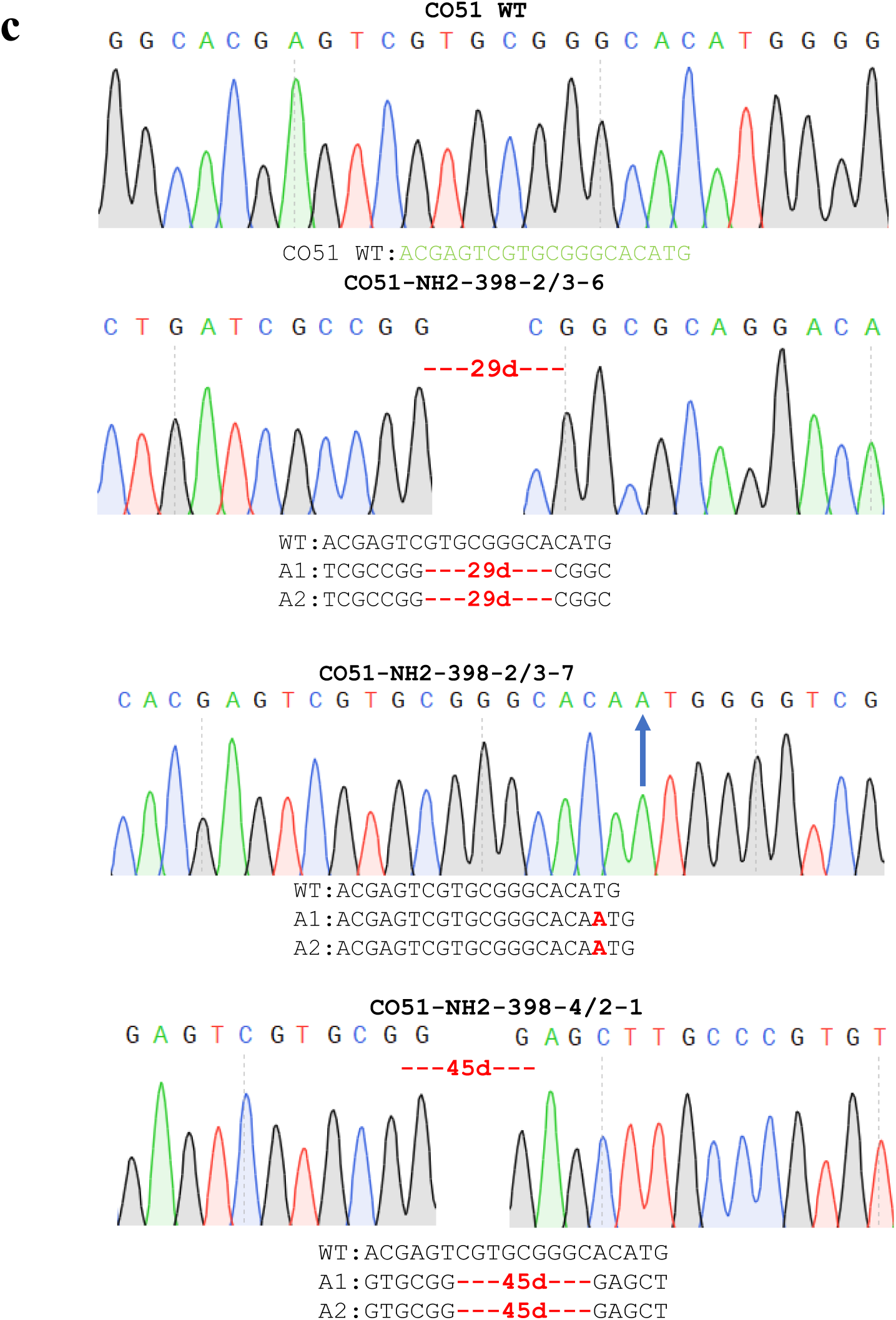
Schematic representation of CRISPR/Cas9-based editing of *OsNH2*. (a) Gene structure and position of sgRNA1 and sgRNA2 at exon 2 along with the PAM sequences. PAM sequences are highlighted in red. The nucleotide positions 398 and 426 indicate gRNA target sites within the exon region. (b) and (c) The chromatogram traces from Sanger sequencing indicate that the homozygous mutants of *OsNH2-398* and *OsNH2-426* show varying mutations compared with WT plants in ASD16 and CO51. The nucleotides highlighted in green represent gRNA sequences. *“d”—deletion; “i”—insertion; “s”—substitution.

#### Off-target analysis and structural impact of *OsNH2* mutants

Sanger sequencing analysis confirmed that no off-target activity was detected in all 7 *OsNH2* mutants in both ASD16 and CO51 backgrounds (Tables S2, S6). Furthermore, due to the high sequence homology observed within the rice NH gene family, a multiple sequence alignment (MSA) of *OsNH2* with other NH genes (*NH1, NH3, NH4,* and *NH5*) revealed a 10 bp imperfect match at the NH2-sgRNA target sites. However, the absence of off-target editing in other NH genes in all mutants indicates that the *OsNH2* mutation alone contributed to the observed phenotypic changes (Table S7).

The WT sequence of *OsNH2* was translated into amino acid sequences, and Interpro analysis revealed that the OsNH2 protein contains several conserved domains similar to those found in the OsNH1 protein, including the BTB/POZ domain, a potassium channel, ankyrin repeats, and the NPR1/NIM1-like defense domain. The alignment of mutant OsNH2 protein sequences with the WT showed partial disruptions in the N-terminal BTB/POZ domain and potassium channel (due to a premature stop codon) and a complete loss of ankyrin repeat and NPR1/NIM1-like defense domains, highlighting the functional impairment of the OsNH2 protein in *OsNH2* mutants (Fig. S1).

### Bioassay of homozygous *OsNH2* mutants (T_2_) against *R. solani* and *Xoo* infection

A bioassay of seven homozygous *OsNH2* mutants (derived from ASD16 and CO51 backgrounds) against *R. solani* was conducted over two generations. A detached leaf bioassay was performed in all the mutants. The lesion area of the leaves was measured at the following time points using ImageJ software: 24, 48, and 72 h post-infection (hpi). At 24 h post-infection, lesion formation began in all *OsNH2* mutants, including the WT counterparts. By 48 hpi, an increase in the number of infection cushions was observed in mutant leaves compared with WT leaves (Fig. S2a–k). WT leaves developed smaller necrotic lesions, whereas mutants showed more extensive necrotic lesions, leading to the yellowing and drying of the entire leaf blade by 72 hpi (Fig. 2a–d). As a result, all *OsNH2* homozygous mutants developed a higher number of lesions compared with WT leaves. Based on the bioassay results, two *OsNH2* mutants from each cultivar that showed the highest disease symptoms were selected for further analysis to investigate the role of *OsNH2* in imparting ShB resistance. In the whole-plant assay, a 3-day-old mycelial plug of *R. solani* was inoculated on both mutants and WT plants. By 3 dpi, lesion formation was observed, with light greyish lesions forming on the sheaths of both WT plants and *OsNH2* mutants. By 7 dpi, *OsNH2* mutants showed higher susceptibility to ShB than their WT counterparts (Fig. 2e–h). They showed increased pathogen spreading and complete drying of infected sheaths, whereas WT plants showed less severe symptoms.

**Fig. 2.**
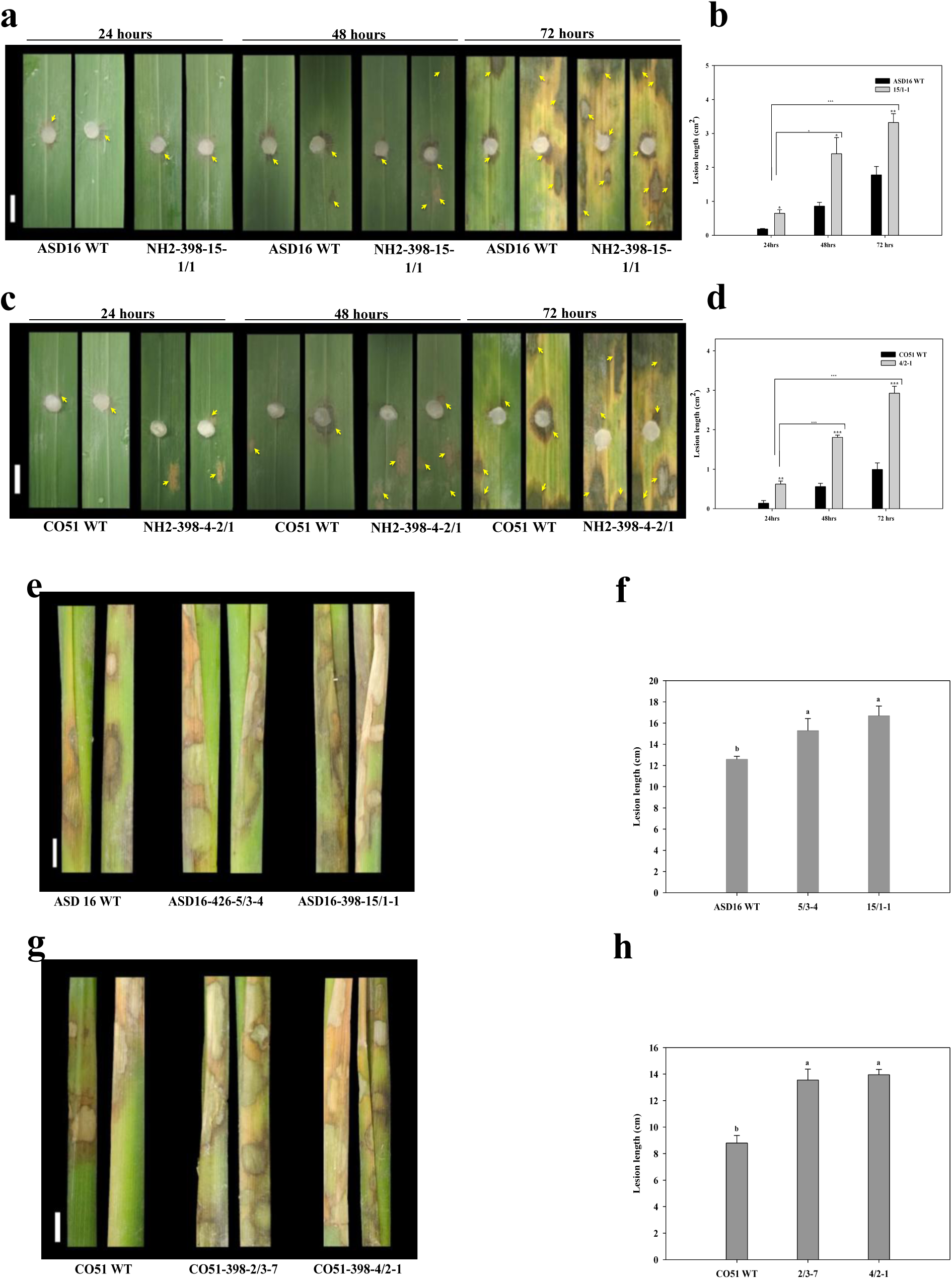
Phenotypic assessment of *R. solani* infection using detached leaf and whole-plant assays. Comparison of lesion formation in WT plants and *OsNH2* mutants in detached leaf bioassay of (a, b) ASD16 and (c, d) CO51. Whole-plant bioassay comparing WT plants and *OsNH2* mutants at 7 dpi and their bar graphs: (e, f) ASD16 and (g, h) CO51. Scale bar = 1cm. Data represent mean ± SE, *n* = 10. Lowercase letters above each bar show significant differences detected by Tukey’s honestly significant difference test. The experiments were carried out in triplicate with similar results, and a representative result is shown. Arrows indicate the necrotic lesion areas.

In addition to the ShB bioassay experiment, *OsNH2* mutants were examined against *Xoo* infection using the leaf clip method. A significant increase in lesion length was observed throughout the infection period in *OsNH2* mutants compared with WT plants, indicating higher susceptibility to *Xoo* infection. After 14 days, the mutants showed more severe disease symptoms than WT plants (Fig. 3). The infection in the mutants progressed to a severe stage, leading to the death of leaves after 5 weeks. This illustrates that *OsNH2* mutants also show higher vulnerability to BLB disease caused by *Xoo*.

**Fig. 3.**
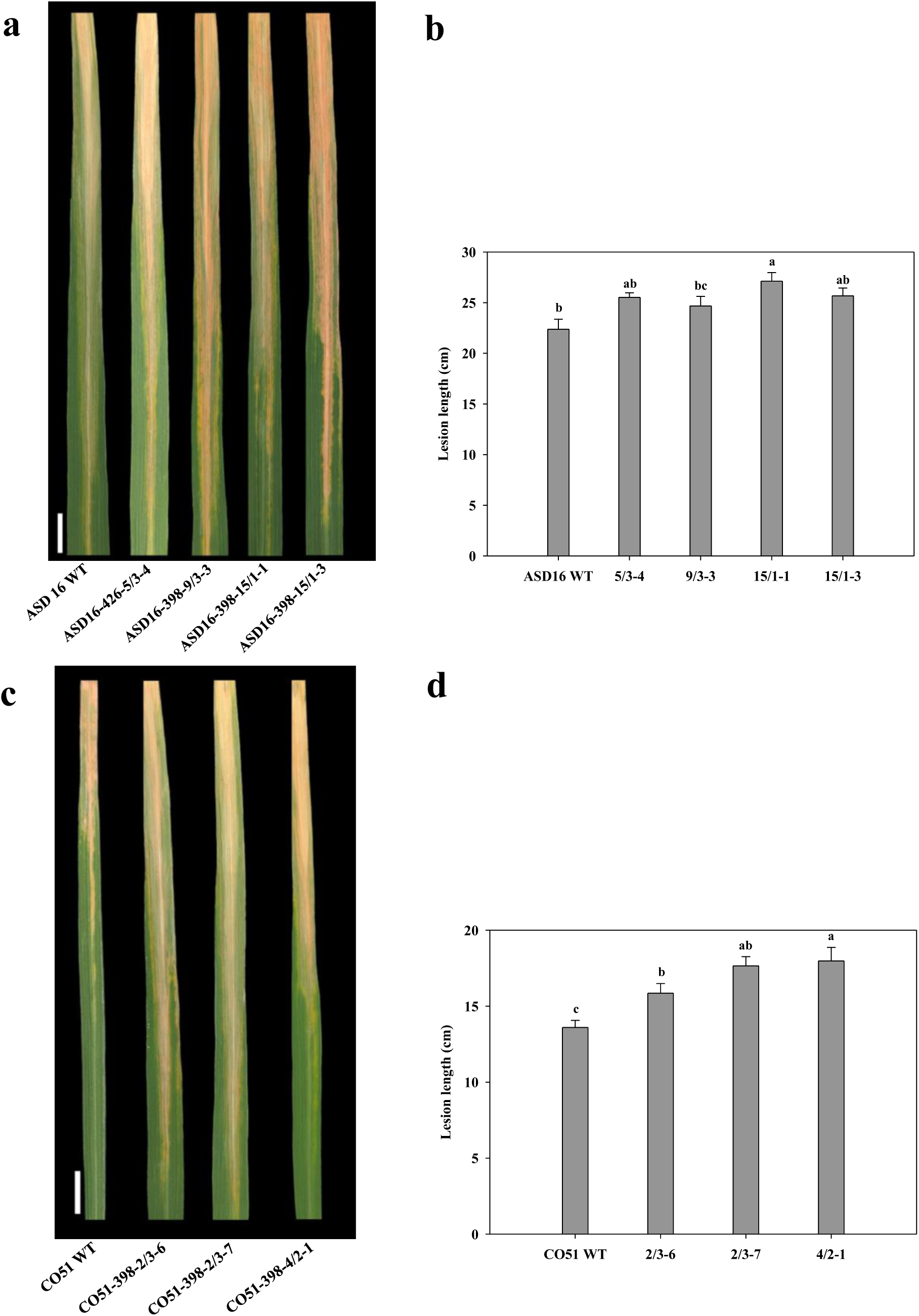
Evaluation of disease symptoms in WT plants and *OsNH2* mutants following *Xoo* infection. (a, b) ASD16 and (c, d) CO51. Scale bar =1cm. Data represent mean ± SE, *n* = 10. Lowercase letters above each bar show significant differences detected by Tukey’s honestly significant difference test. The experiments were carried out in triplicate with similar results, and a representative result is shown.

#### ROS accumulation and SEM analysis in *OsNH2* mutant leaves infected with *R. solani*

Superoxide, a major reactive oxygen species (ROS), was detected in *R. solani*-infected rice leaves using NBT staining at 48 hpi in two selected *OsNH2* mutants from ASD16 and CO51 cultivars. Superoxide accumulation resulted in higher blue polymerization in *OsNH2* mutants than in WT leaves. Among mutants, ASD16-398-15/1-1 and CO51-398-4/2-1 showed the highest ROS accumulation, whereas WT plants showed the least amount of oxidation product (Fig. S3). SEM analysis at 48 hpi visualized dense, coalesced mycelial networks of *R. solani* on both *OsNH2* mutant and WT leaves. Mutants also exhibited higher hyphal density, greater hyphal width, and increased branching compared to WT plants (Fig. S4). Notably, infection cushions formed only in mutants (Fig. S4e–f), while WT leaves lacked these structures. These observations suggest that the *OsNH2* mutation enhances fungal colonization and increases susceptibility to *R. solani*.

### Defense-related gene expression in *OsNH2* mutants upon *R. solani* infection

The mRNA expression levels of key defense-related genes, including PR genes and transcription factors from the WRKY and TGA families, were analyzed in *OsNH2* mutants and WT plants in both cultivars. Expression analysis of TGA and WRKY transcription factors showed significantly reduced levels of *OsWRKY4*, *OsWRKY45, OsTGA2*, and *OsTGA3* in *OsNH2* mutants compared to WT at 24 and 72 hpi. However, *OsWRKY80* showed differential expression only at 72 hpi, while *OsTGA5* expression was not altered in response to *R. solani* infection (Figs. 4 and 5). Similarly, the expression of defense-related genes *OsPR1*, *OsPR3*, *OsPR5*, and *OsPR10* (P>0.05) was significantly lower in mutants at both time points, with reduced basal expression as well (Fig. S5). Taken together, the reduced expression of these transcription factors and PR genes suggests suppression of defense pathways in *OsNH2* mutants.

**Fig. 4.**
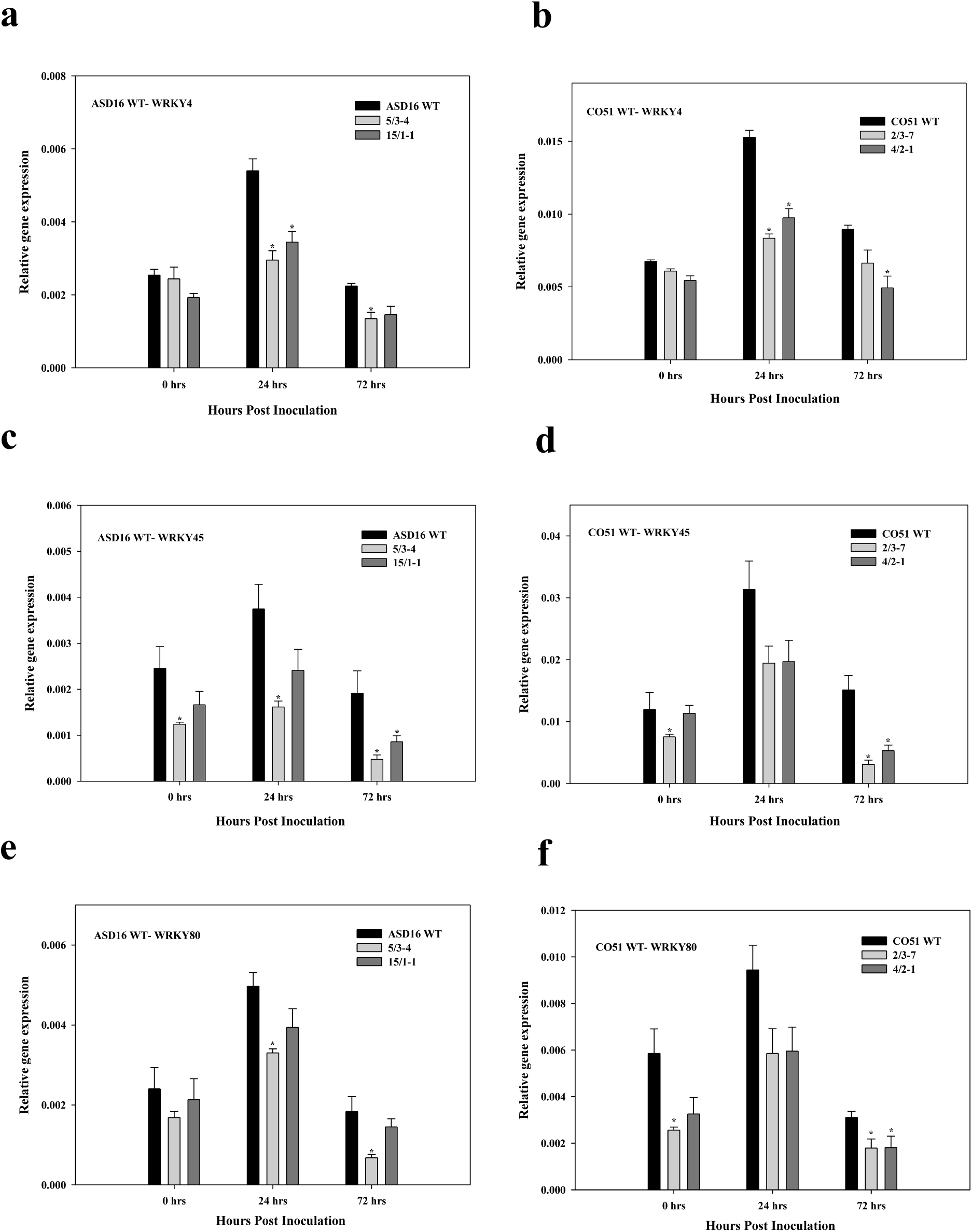
Expression analysis of WRKY transcription factors in WT plants and *OsNH2* mutants. (a, b) *OsWRKY4*, (c, d) *OsWRKY45*, and (e, f) *OsWRKY80*. Data represent mean ± SE. Asterisks indicate significant differences (*P < 0.05 Student’s t-test) between different time intervals. The experiment was performed in duplicate with consistent results.

**Fig. 5.**
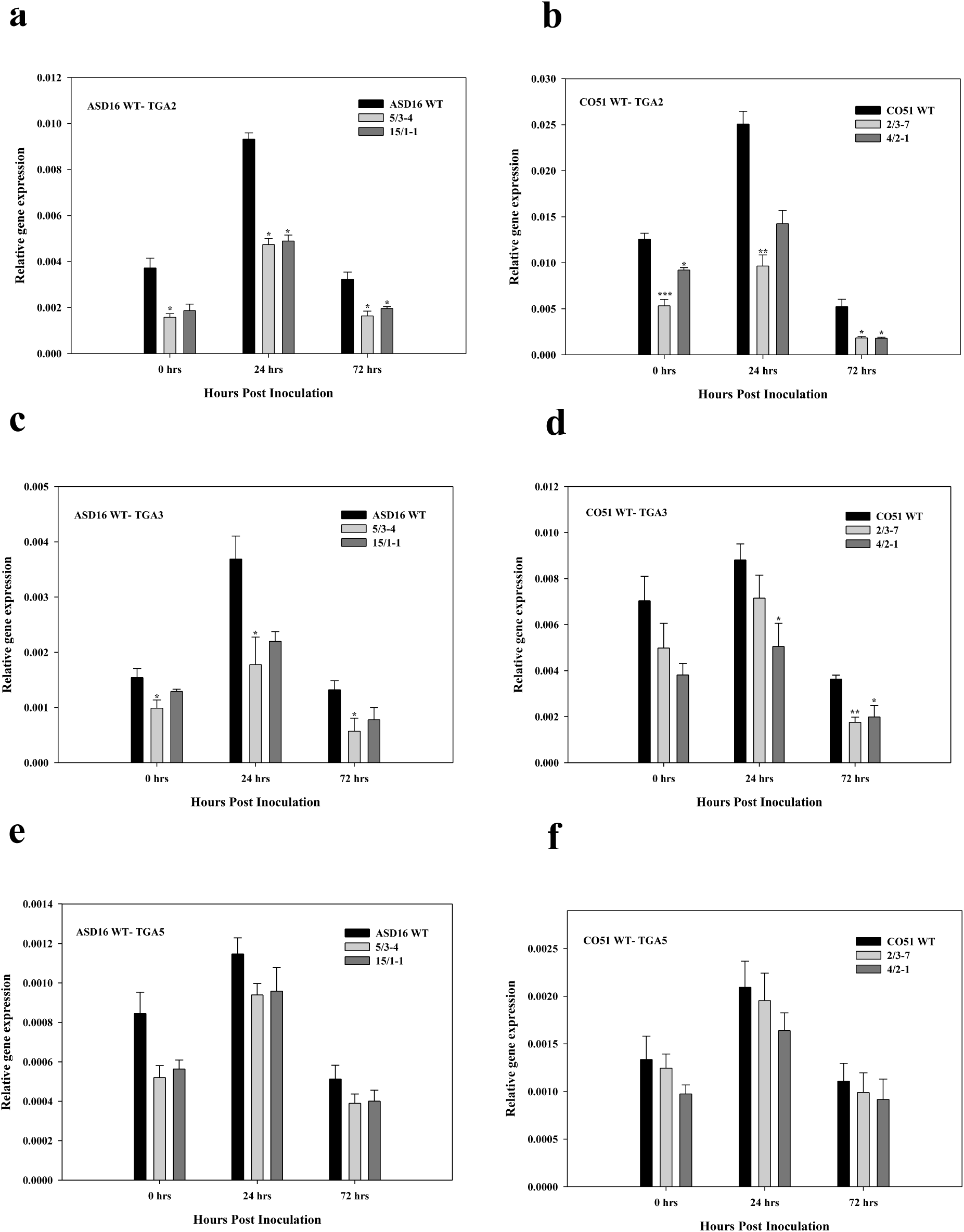
Expression analysis of TGA transcription factors in WT plants and *OsNH2* mutants. (a, b) *OsTGA2*, (c, d) *OsTGA3*, and (e, f) *OsTGA5*. Data represent mean ± SE. Asterisks indicate significant differences (*P < 0.05, **P < 0.01, ***P < 0.001, Student’s t-test) between different time intervals. The experiment was carried out in duplicate with reproducible results.

### Analysis of the expression levels of *OsNH1* and *OsNH3* in *OsNH2* mutants

To further explore the regulatory role of *OsNH2*, we examined the expression of *OsNH1* and *OsNH3* in the mutants. Both *OsNH1/3* genes showed significantly lower expression in mutants compared to WT plants, indicating that *OsNH2* is required for their proper expression (Fig. 6a–d). Additionally, *R. solani* infection significantly induced *OsNH2* expression in WT plants, while less induction was observed in the mutants (Fig. 6e–f).

**Fig. 6.**
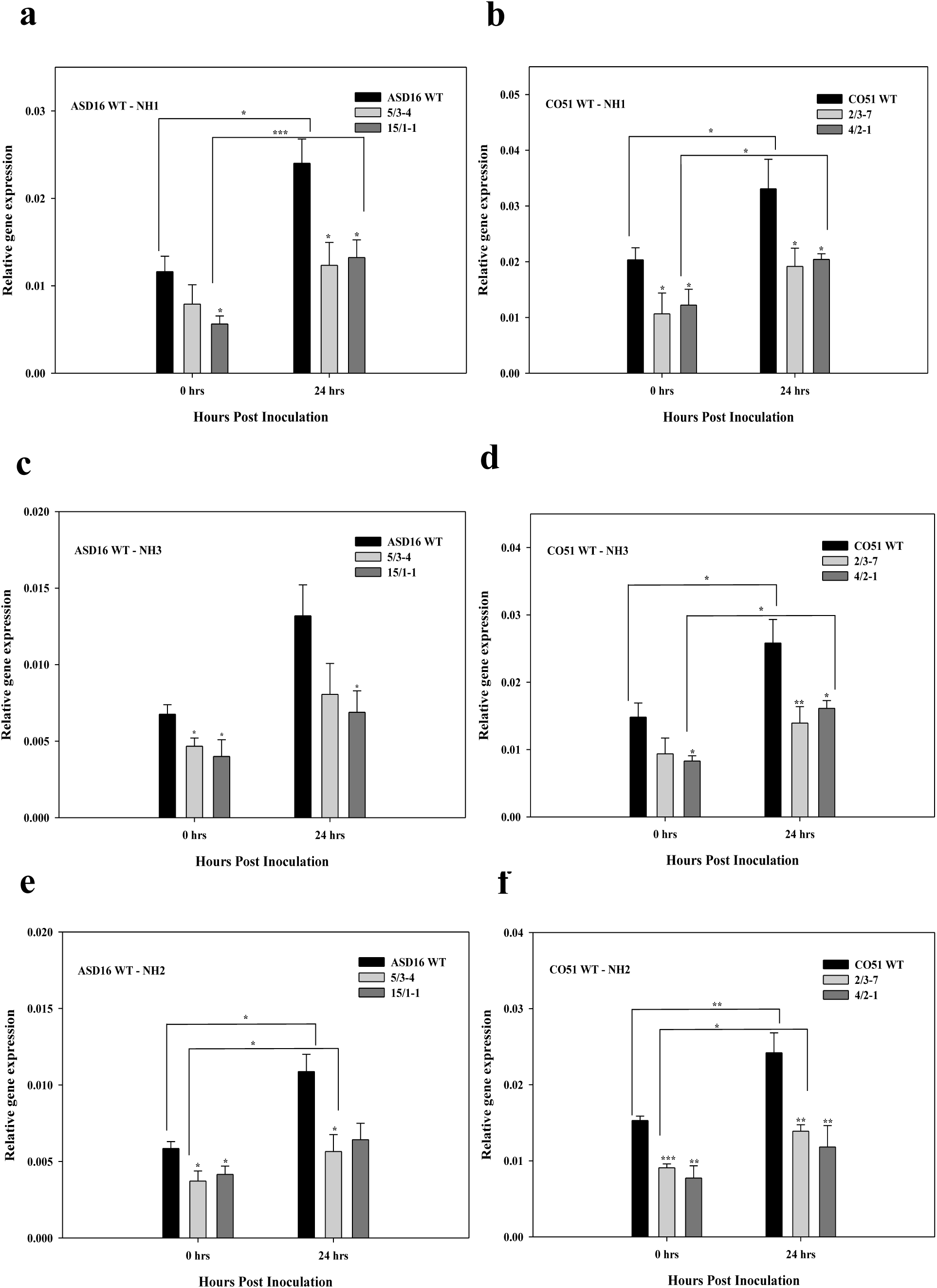
***OsNH1/3* expression in WT plants and *OsNH2* mutants**. (a, b) *OsNH1*, (c, d) *OsNH3*, and (e, f) *OsNH2*. Data represent mean ± SE. Asterisks indicate a significant difference (*P < 0.05, **P< 0.01, ***P< 0.001 Student’s t-test) between different time intervals. The experiment was carried out in duplicate with similar trends among each other.

#### Effect of exogenous application of phytohormones on *OsNH2* mutants during *R. solani* infection

To investigate whether exogenous phytohormone application enhances the immune response against ShB disease, WT and *OsNH2* mutants were sprayed with 1 mM of SA, ET, and JA. After 24 h of spray, leaves were detached and inoculated with *R. solani*. At 48 hpi, SA-treated WT and SA-treated *OsNH2* mutants exhibited significantly reduced symptoms compared to their respective controls (Fig. 7 and Fig. S6). However, the extent of lesion reduction in mutants was significantly lower than in WT plants, underscoring the pivotal role of *OsNH2* in SA-mediated defense signaling. ET application did not alter disease severity, whereas JA treatment moderately enhanced susceptibility in this pathosystem (data not shown).

**Fig. 7.**
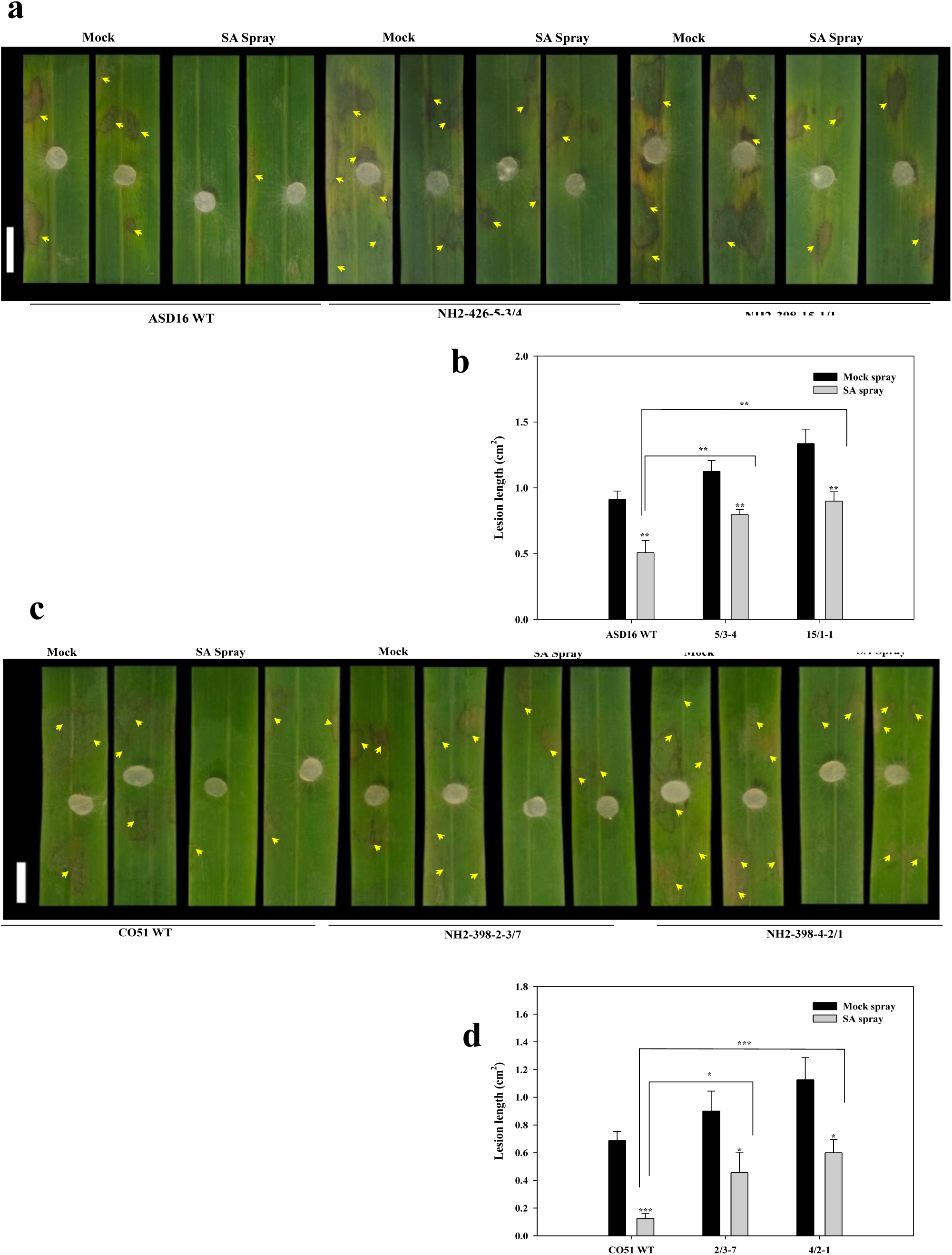
Effects of exogenous SA application on WT plants and *OsNH2* mutants 24 h after spray, followed by detached leaf assay. (a, b) ASD16 (c, d) CO51. Mock indicates water spray, SA—Salicylic acid (1 mM). The lesion area was measured using ImageJ software after the 48 hpi and plotted on a graph. Scale bar =1cm. Data represent mean ± SE, *n* = 20. Asterisks indicate a significant difference (*P < 0.05, **P< 0.01, ***P< 0.001 Student’s t-test). The experiments were carried out in triplicate, showing consistent results. Arrows indicate the necrotic lesion areas.

Further, endogenous SA levels were quantified in ASD16 and CO51 WT plants and *OsNH2* mutants under water spray, SA spray, and *R. solani* inoculation treatments. Before infection, SA content did not differ between WT and mutants in both cultivars. However, after the SA spray, WT plants accumulated higher SA than *OsNH2* mutants. Upon *R. solani* infection, SA levels increased in both water- and SA-sprayed WT plants, while mutants showed a significantly smaller increase compared to WT plants, demonstrating that *OsNH2* is required for maintaining elevated SA levels during pathogen challenge (Fig. 8). The expression levels of *OsNH1* and *OsNH3* were analyzed in SA-treated ASD16 WT and SA-treated *OsNH2* mutants at 24 hpi. These experiments were performed exclusively in the ShB susceptible ASD16 cultivar, as SA spray was observed to confer a comparable level of symptom reduction in both ASD16 and CO51 cultivars. SA treatment significantly upregulated the expression of *OsNH1*/*3* in ASD16 WT plants. Notably, *OsNH2* mutants also responded to SA spray, showing a significant increase in the expression of *OsNH1*/*3* compared to their water-treated mutants, although the levels remained lower than those observed in ASD16 WT plants (Fig. S7a and b). This suggests that *OsNH2* is required for the full activation of *OsNH1* and *OsNH3.* Additionally, SA treatment significantly induced *OsNH2* expression in WT plants compared to water-treated controls (Fig. S7c).

**Fig. 8.**
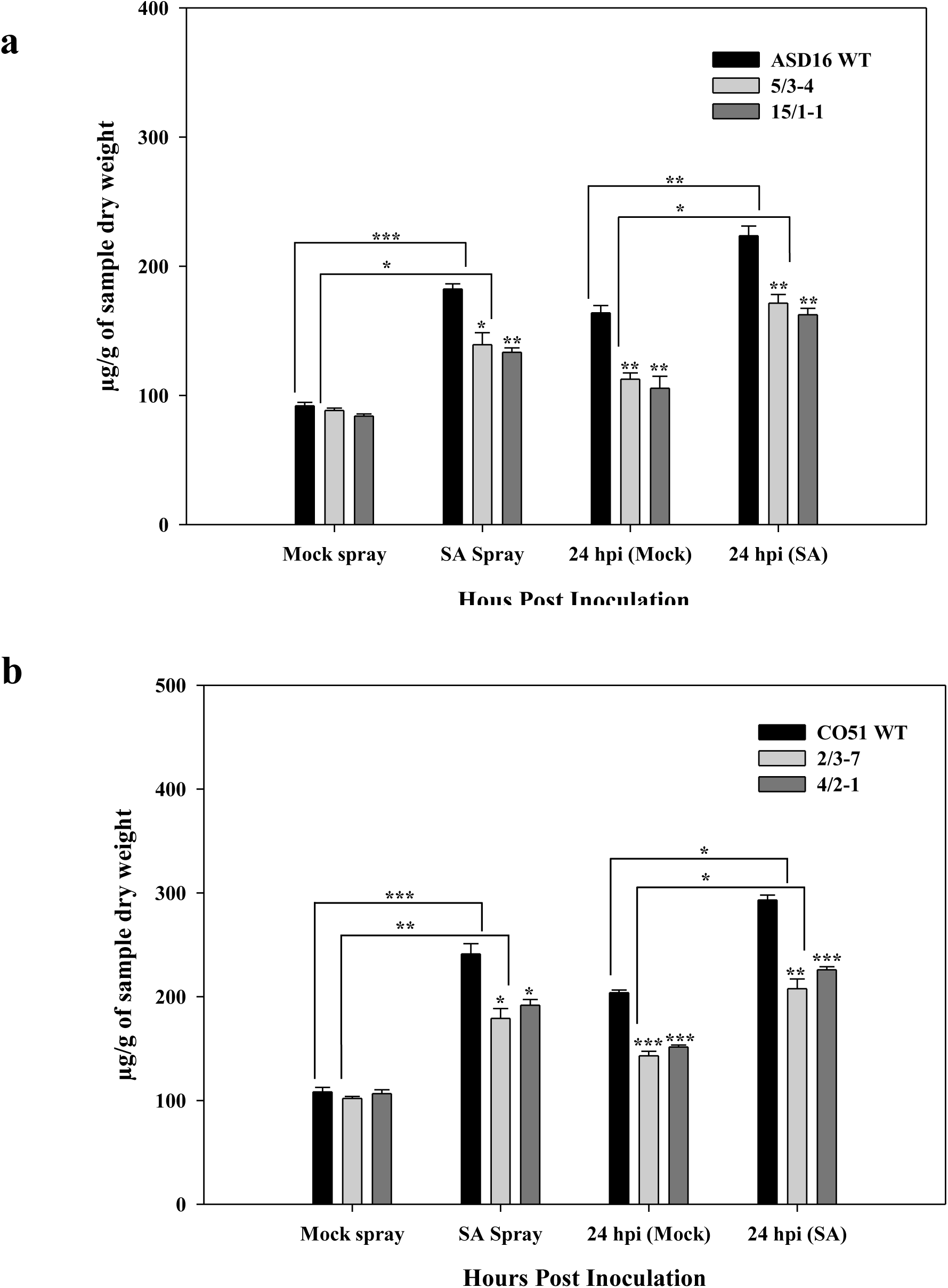
Quantification of SA content in WT plants and *OsNH2* mutants. (a) ASD16 and (b) CO51. Endogenous SA content was measured in WT plants and mutants under mock and SA treatments, before and after *R. solani* infection at 24 hpi, and are expressed as µg/g of dry weight. Data represent mean ± SE. Asterisks indicate significant differences (*P < 0.05, **P < 0.01, ***P < 0.001, Student’s t-test). The experiment was repeated three times with consistent results.

## Discussion

*OsNH2* mutants developed using CRISPR/Cas9 technology exhibited increased susceptibility to ShB and BLB. The expression of defense-related genes, including WRKYs, TGAs, PR genes, and NPR1 homologs such as *OsNH1* and *OsNH3*, was significantly reduced in the mutants. The mutants also showed enhanced ROS accumulation, increased hyphal density of *R. solani*, and reduced SA content. Exogenous SA application partially restored resistance in the mutants against *R. solani* infection.

### Susceptibility analysis of *OsNH2* mutants to *R. solani* and *Xoo* using bioassay studies

The phenotypic assessment of *OsNH2* mutants in ASD16 and CO51 backgrounds showed increased susceptibility to *R. solani*. In particular, even the moderately resistant cultivar CO51 became susceptible to ShB infection due to the *OsNH2* mutation, underscoring the critical role of this gene in defense (Fig. S2). Similarly, *OsNH2* mutants showed increased susceptibility to *Xoo* infection compared to WT plants (Fig. 3). This finding is consistent with earlier studies where knockdown of *OsNPR1/OsNH1* led to increased susceptibility to *Xoo* compared to wild-type (Yuan et al., 2007). Previous studies have reported that knockout or knockdown of rice signal transduction pathway genes, *PP2A-1*, *CIPK31*, and *MDPK*, results in increased susceptibility to the ShB (Lin et al., 2021; Cui et al., 2022; Chen et al., 2023). Collectively, these results indicate that *OsNH2* also plays a positive regulatory role in defense, similar to *OsNH1*, against both fungal and bacterial pathogens.

#### Functional domain analysis in OsNH2 Protein

*AtNPR1* and its rice homolog, *OsNH1* exhibit significant similarities and play crucial roles in plant immunity by regulating SA signaling and activating defense-related genes during pathogen infections (Cao et al., 1994; Zhang et al., 1999). Pairwise sequence identity revealed that OsNH2 shares the highest similarity with OsNH1, followed by OsNH3, suggesting a functional relationship among the NPR1-like proteins in rice (Fig. S8). In addition, the OsNH1 protein is characterized by multiple functional domains, including the BTB/POZ, potassium channel, ankyrin repeat, and NPR1/NIM1-like defense protein, which contribute to diverse roles in plant–pathogen interactions (Aravind and Koonin, 1999; Li et al., 2006). The OsNH2 protein displays domain similarities to AtNPR1, OsNH1, and OsNH3, suggesting a potential positive regulatory function in rice immunity (Fig. S9). The *OsNH2* mutants generated by targeting the initial exons resulted in truncated proteins lacking all these functional domains (Fig. S1). Previous studies have shown that various domains of NPR proteins—BTB/POZ domain, potassium channel-like domain structurally related to POZ, ankyrin repeats, and NPR1/NIM1-like domain—are essential for their interaction with TGA transcription factors, activation of immune signaling, and expression of PR genes (Li et al., 2006; Aravind and Koonin, 1999; Maier et al., 2011; Boyle et al., 2009). The absence of any of these domains in NPR proteins impairs hormonal signaling in the SA and JA/ET pathways, thereby compromising the plant’s immune response (Spoel et al., 2003). Notably, mutation in the ankyrin repeat domain of the Arabidopsis *npr1* mutant allele completely lost responsiveness to SAR, reduced PR gene expression, and increased susceptibility to fungal and bacterial pathogens (Cao et al., 1997). Accordingly, the *OsNH2* frameshift mutants generated in the study, which lack all functional domains, are likely to be nonfunctional and exhibit increased susceptibility to *R. solani* and *Xoo*, demonstrating that *OsNH2* functions similarly to *AtNPR1/OsNH1,* but with a less prominent role in rice defense than *OsNH1/3*.

In addition, the in-frame mutant CO51 4/2-1 (derived from the moderately ShB-resistant cultivar), which lacks amino acids 121–135 within the potassium channel/BTB domain, exhibited ShB disease severity similar to that of the *OsNH2* knockout mutants. Similarly, another in-frame mutant, ASD16 15/1-3, lacking amino acids 134–151 in the same domain, also showed comparable disease severity, further supporting the notion that an intact *OsNH2* protein is essential for enhanced defense (Fig. S2, Table S8).

#### Increased ROS accumulation and hyphal width in *OsNH2* mutants in response to *R. solani* infection

Chloroplasts are a significant source of ROS, which act as signaling molecules in plant defense (Hu et al., 2021). ROS restrict biotrophic pathogens by inducing programmed cell death but can promote necrotrophic pathogens, such as *R. solani*, that thrive on dead tissue (Torres et al., 2006; Zhang et al., 2020). NBT staining techniques showed that superoxide typically increases following *R*. *solani* infection (Foley et al., 2016). The *OsNH2* mutants exhibited higher superoxide levels (Fig. S3), correlating with enhanced disease susceptibility to *R. solani* and *Xoo*. This finding aligns with an earlier report demonstrating that *OsGSTU5* knockdown lines accumulated higher ROS levels and showed increased susceptibility to *R. solani* (Tiwari et al., 2020).

SEM analysis further demonstrated increased hyphal width and enhanced infection cushion formation in *OsNH2* mutants at 48 hpi with *R. solani*, underscoring the importance of intact *OsNH2* in rice defense. Previous SEM studies on resistant and susceptible rice varieties infected with *R. solani* have demonstrated the utility of this technique in evaluating fungal infection levels (Cao et al., 2022). In our analysis, ASD16 WT displayed dense hyphal growth, indicating higher susceptibility, whereas CO51 showed sparse hyphal growth, reflecting moderate ShB resistance. The lesion length measurement in bioassay also correlates with SEM data, validating the susceptibility in ASD16 and moderate resistance in CO51. Notably, *OsNH2* mutants in both rice cultivars displayed greater hyphal width than their respective WT plants, highlighting the role of a fully functional *OsNH2* in defense, similar to *OsNH1/3*. Surprisingly, SEM analysis confirmed that even CO51—a moderately ShB-resistant variety—became susceptible to *OsNH2* mutation (Fig. S4). Similar results were observed in an in-frame mutant of CO51, further reinforcing the critical role of *OsNH2* in rice ShB defense.

#### *OsNH2* is essential for rice defense and maintaining *OsNH1/3* expression

*R. solani* infection significantly increased *OsNH2* expression in WT plants, suggesting its involvement in defense responses. In contrast, mutants showed reduced expression of *OsNH2*, indicating a potential loss of function in immunity (Fig. 6e–f). The *OsNH1* and *OsHN3* expression levels were also lower in *OsNH2* mutants than in WT plants, confirming that *OsNH2* positively regulates the expression of *OsNH1/3.* Previous studies have demonstrated that overexpression of *AtNPR1, OsNPR1,* and *OsNPR3*.3 in rice conferred disease resistance to ShB and *Xoo* (Molla et al., 2016; Jiang et al., 2020; Dai et al., 2023), underscoring the critical role of NPR-family genes in defense. Bai et al., (2011) reported that the overexpression of *OsNH3* in rice conferred enhanced resistance to *Xoo*, accompanied by a 3.5-fold increase in *OsNH1* expression following benzothiadiazole treatment. Similarly, their microarray data indicated that overexpression of *OsNH1* resulted in a 1.6-fold upregulation of *NH3* expression upon benzothiadiazole spray. In a recent study, Dai et al., (2023) demonstrated that *OsNPR1*-overexpressing rice lines displayed resistance to BLB, along with increased expression of *OsNPR3* upon *Xoo* infection. These studies establish a positive regulatory interplay between NPR1 and NPR3 during *Xoo* infection. However, the interactions among the NH1/2/3 gene family in regulating defense signaling networks remain unclear. This study proves the importance of *OsNH2* in rice defense and its essential role in inducing the expression of *OsNH1/3* to mediate SA-mediated defense.

Further, the higher expression levels of *OsNH1/3* in CO51 WT compared to ASD16 WT emphasize their role in rice defense (Fig. 6a–d), while the marked downregulation of *OsNH1/*3 in *OsNH2* mutants appears to be a key factor contributing to the increased susceptibility to ShB. These findings highlight the complex regulatory network of NH genes in rice defense against various pathogens, with *OsNH1/2/3* each contributing uniquely to modulating disease resistance.

#### Loss of *OsNH2* disrupts the SA-mediated defense network

In *OsNH2* mutants, key transcription factors—*OsWRKY4*, *OsWRKY45*, *OsWRKY80*, *OsTGA2*, and *OsTGA3*—were downregulated, which are known to regulate SA-dependent defense pathways. Previous studies have shown that WRKY and TGA transcription factors control the expression of *PR* genes during *R. solani* infection (Kouzai et al., 2016; Peng et al., 2016; Zhou et al., 2000; Backer et al., 2019). Overexpression of *OsNPR1/3* in rice significantly upregulated WRKY and PR genes, including WRKY45, PR1a, PR5, and PR10a, during *Xoo* infection (Bai et al., 2011; Dai et al., 2023). In our study, the downregulation of WRKY45 and PR genes (PR1, PR5, PR10) in *OsNH2* mutants correlates with increased susceptibility to *Xoo* and ShB.

A coordinated interaction between the NH family genes and TGAs has been reported to regulate the SA signaling cascade (Moon et al., 2018; Chern et al., 2014). STRING-based protein interaction analysis revealed strong associations of NH2 with NH1/3 and TGA2.2/2.3, moderate interaction with NH5, weak interaction with NH4, and no interaction with TGA3 (Fig. S10). The NH5—previously uncharacterized in plant defense (Jun et al., 2010; Chern et al., 2021)—showed strong to moderate protein interactions with NH1/2/3, and a recent study demonstrated that a novel variant, *OsNH5N16*, confers resistance to BLB and Bakanae disease in rice (Son et al., 2021). STRING analysis showed that OsTGA2.2/2.3 interact strongly with NH2 and NH1/3, supporting their involvement in defense signaling. In contrast, no interaction was observed with *OsTGA3,* which aligns with earlier findings that overexpression of *OsTGA3* did not contribute to defense (Moon et al., 2018). NH2 has very weak interactions with NH4, consistent with previous findings that its overexpression fails to enhance disease resistance (Bai et al., 2011). Overall, this *in silico* study supports that NH2 functions in coordination with NH1/3 and possibly NH5 to regulate defense signaling rather than acting independently. In *OsNH2* mutants, disruption of this interconnected network impairs SA-mediated signaling, resulting in increased susceptibility to *R. solani*.

SA is synthesized through two main pathways—the isochorismate synthase (ICS) pathway and the phenylalanine ammonia-lyase (PAL) pathway (Lefevere et al., 2020). In *OsNH2* mutants, the SA biosynthetic genes *OsPAL* (p<0.05) and *OsICS1* (p>0.05) were not significantly upregulated compared to the WT (Fig. S11), correlating with lower SA content in the mutants following *R. solani* infection. These genes are typically regulated by WRKY and TGA transcription factors upon NPR1/NH1 activation, thereby establishing a positive feedback loop (Shah, 2003; Mishra et al., 2024). In *OsNH2* mutants, SA biosynthesis genes were less upregulated during *R. solani* infection due to reduced expression of WRKY and TGA transcription factors, which are known to activate defense signaling. As infection progressed, the mutants failed to enhance SA production to levels observed in the WT, leading to a compromised immune defense. After *R. solani* infection, both mock- and SA-treated WT plants and mutants differed significantly (P < 0.05). However, SA-sprayed mutants did not show a significant increase in SA content compared with mock-treated WT plants, indicating only partial uptake/response of exogenous SA in the absence of *OsNH2*, thereby underscoring its critical role in regulating SA-mediated defense. Notably, CO51 WT and its mutants exhibited significantly higher basal and pathogen-induced SA levels than ASD16, which may contribute to their enhanced resistance against ShB infection (Fig. 8).

Additionally, docking analysis confirmed that SA binds to OsNH1 (−5.1 kcal mol⁻¹), OsNH2 (−6.1 kcal mol⁻¹), and OsNH3 (−5.8 kcal mol⁻¹), indicating potential interactions with these proteins. Notably, OsNH2 showed the strongest binding affinity and most stable conformation, suggesting a key role in SA signaling (Fig. S12). In both the knockout and in-frame mutants, disruption or loss of the SA-binding domain (C-terminal NPR1/NIM1-like defense) likely impaired SA binding, thereby weakening the overall defense response to *R. solani* infection in rice compared to the WT plants.

#### The function of *OsNH2* in phytohormone-based immunity against rice ShB disease

Understanding the role of phytohormones in regulating plant defense signaling is critical for developing effective strategies to enhance rice resistance to ShB disease (Bari and Jones, 2009; Kouzai et al., 2018). In this study, SA spray effectively reduced ShB disease symptoms (p<0.05) (Fig. 7) in WT and *OsNH2* mutants. Exogenous SA application is known to enhance plant defense by modulating the SA signaling pathway against *R. solani* (Kouzai et al., 2018). Following SA treatment in the WT plants, reduced *R. solani* infection was observed, along with enhanced expression of *OsNH2,* including *OsNH1/3*. The *OsNH2* mutants treated with SA also exhibited reduced ShB lesion sizes and increased expression of *OsNH1/3*, although neither reached the levels observed in WT plants (Fig. 7 and, Fig. S7). These results suggest that while exogenous SA can partially restore the defense response in *OsNH2* mutants, full restoration of the immune function requires a functional NH2 gene.

Thus, the NH2 protein appears to act in coordination with NH1/3 to positively regulate the SA-mediated signaling pathway to maintain effective immunity against *R. solani*.

Overall, these results suggest that *OsNH2* is not acting in isolation but is part of an integrated immune signaling hub, coordinating NPR1 homologs and associated transcription factors to ensure robust defense gene activation in rice. Disruption of any component within this network compromises the plant’s ability to mount an effective defense. However, further studies are required to delineate the precise molecular mechanisms and hierarchical roles of these interacting components in the defense pathway.

## Statements

**Acknowledgments:** The authors express their sincere gratitude to the Department of Plant Biotechnology, Centre for Plant Molecular Biology and Biotechnology, Tamil Nadu Agricultural University, Coimbatore, for providing research facilities. The authors also thank the Central University of Kerala, Kasaragod, for providing the *Xoo* strain to conduct bioassays.

**Author Contributions:** PV, AS, and SV conceptualized and designed the research study. PV and AS conducted the formal analysis, collected the data, and prepared the original draft. DJ and SV validated the statistical analysis. KK, DS, VP, LA, EK, CG, DJ, and SV critically reviewed and edited the manuscript. SV carried out overall supervision and acquired funding for the research. All authors approve the final version of the manuscript.

**Funding:** This work was supported by grants from the Department of Biotechnology, New Delhi, India (BT/PR40456/AGIII/103/1248/2020) and Tamil Nadu Agricultural University, Coimbatore, India.

**Data Availability:** The original contributions presented in the study are included in the article/supplementary material. Further inquiries can be directed to the corresponding author.

## Declarations

**Conflict of interest:** The authors report no conflict of interest.

---

**Fig. S1.**
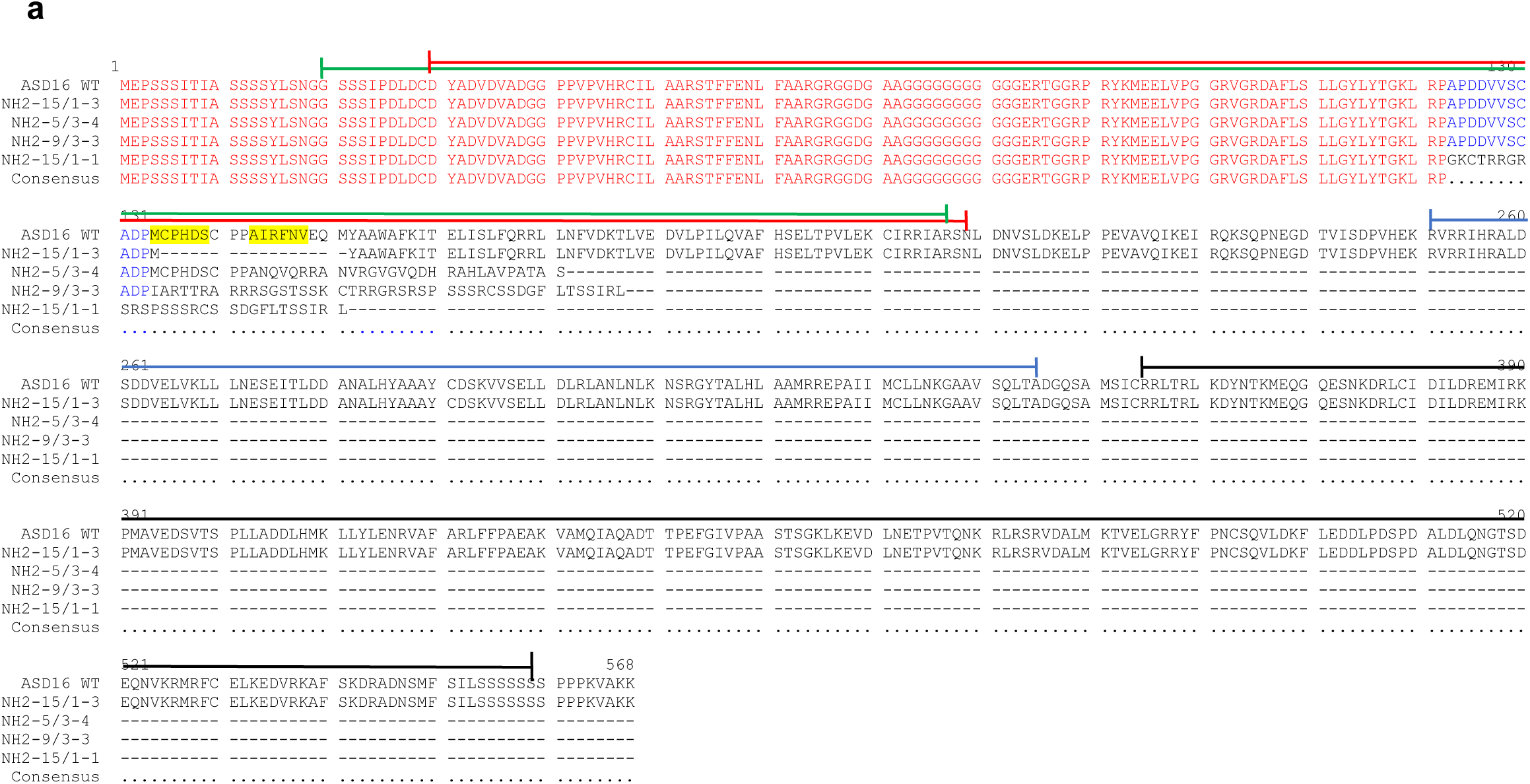
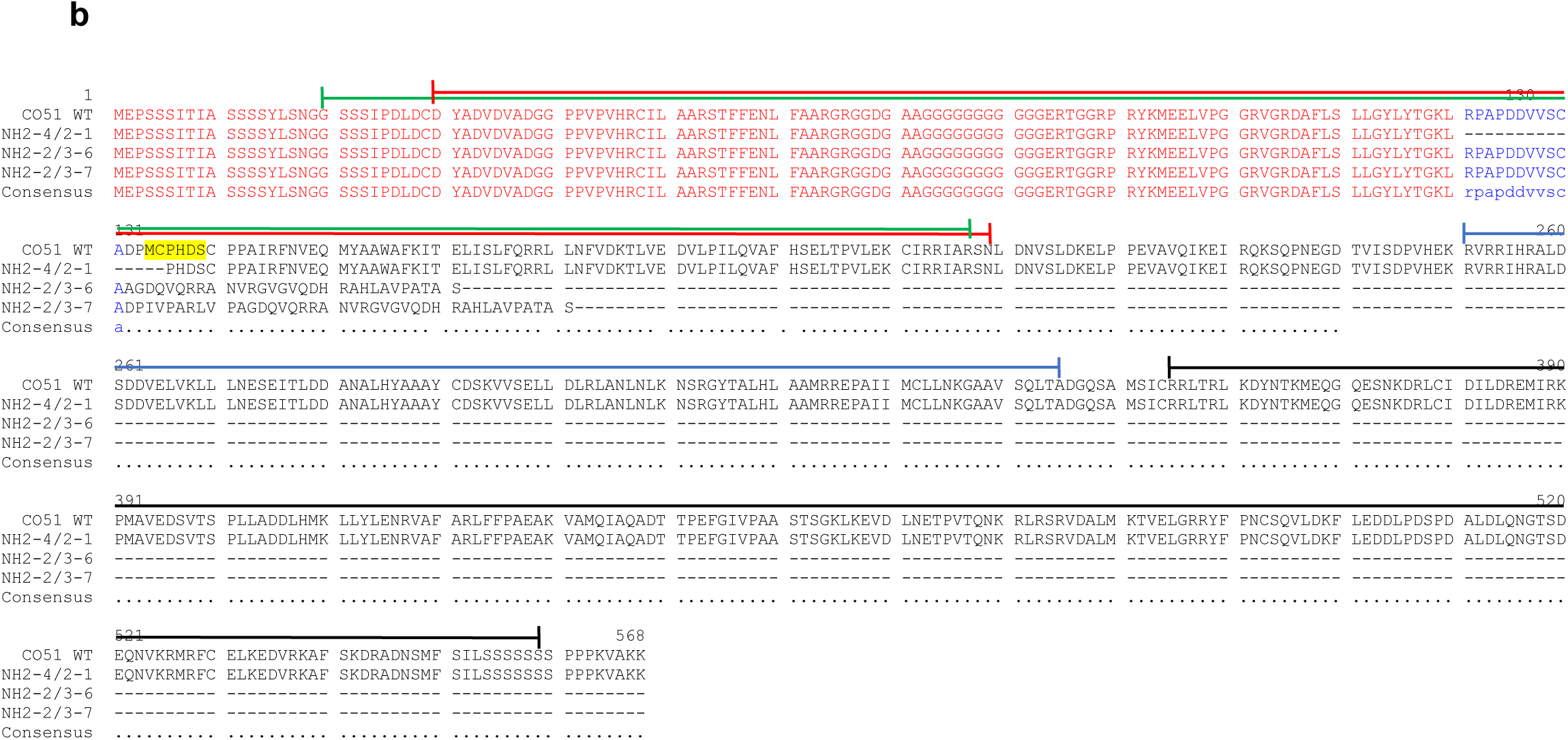
Multiple sequence alignment of WT and *OsNH2* mutant proteins, highlighting key functional domains. The alignment features are color-coded as follows: (a, b) In ASD16 and CO51 mutants, **Green** highlights the potassium channel Kv1.1, **Red** indicates the BTB/POZ domain, **Blue** represents the ankyrin repeats, and **Black** denotes the NPR1/NIM1-like defense protein C-terminal region. In both the cultivars, a premature stop codon results in a truncated protein with reduced amino acid length, causing alterations in key functional domains such as the BTB/POZ domain and the potassium channel domain, while leading to the loss or alleviation of ankyrin repeats and NPR1/NIM1-like defense protein domains. **Yellow** indicates the sgRNA-targeted region in the protein sequence.

**Fig. S2.**
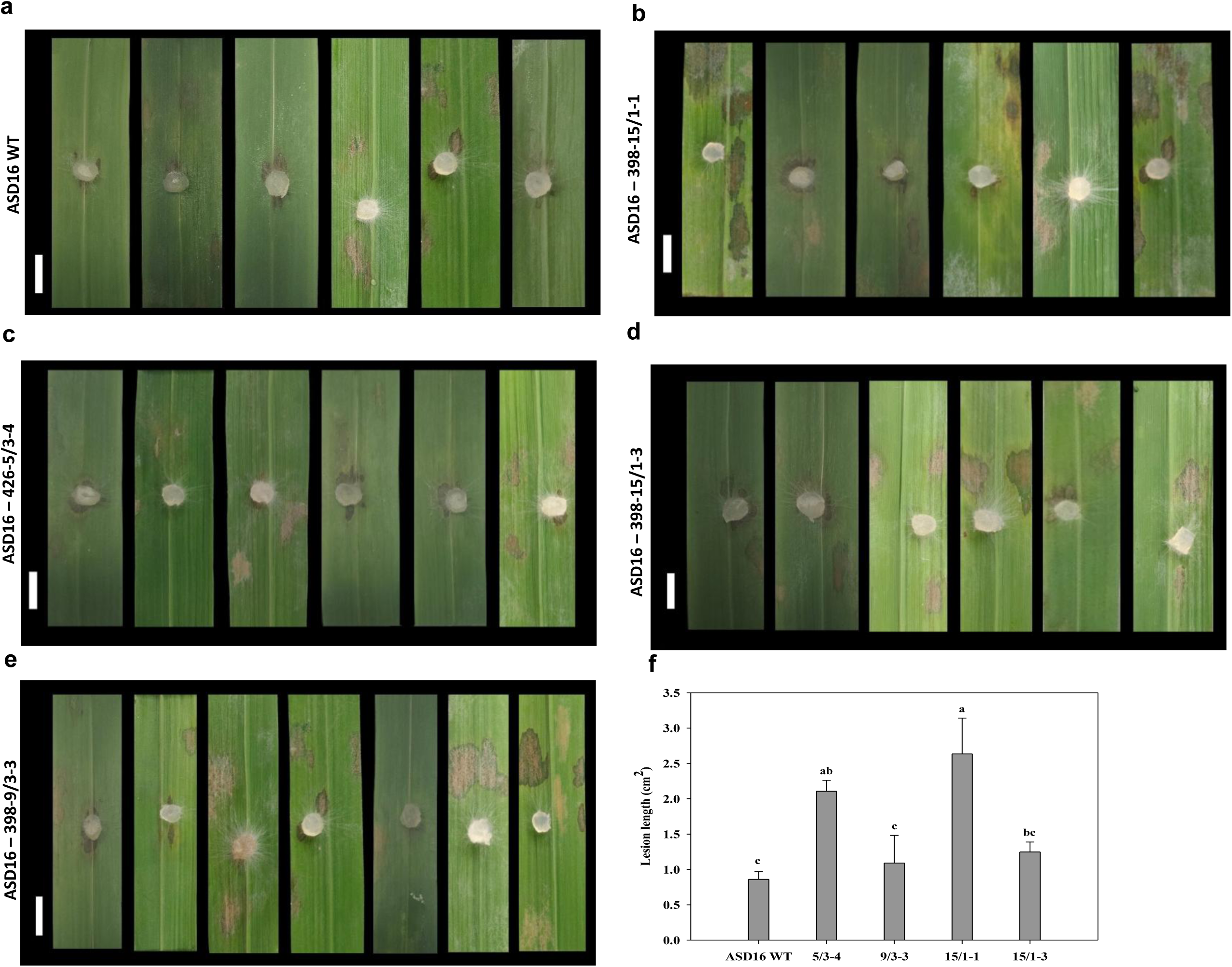
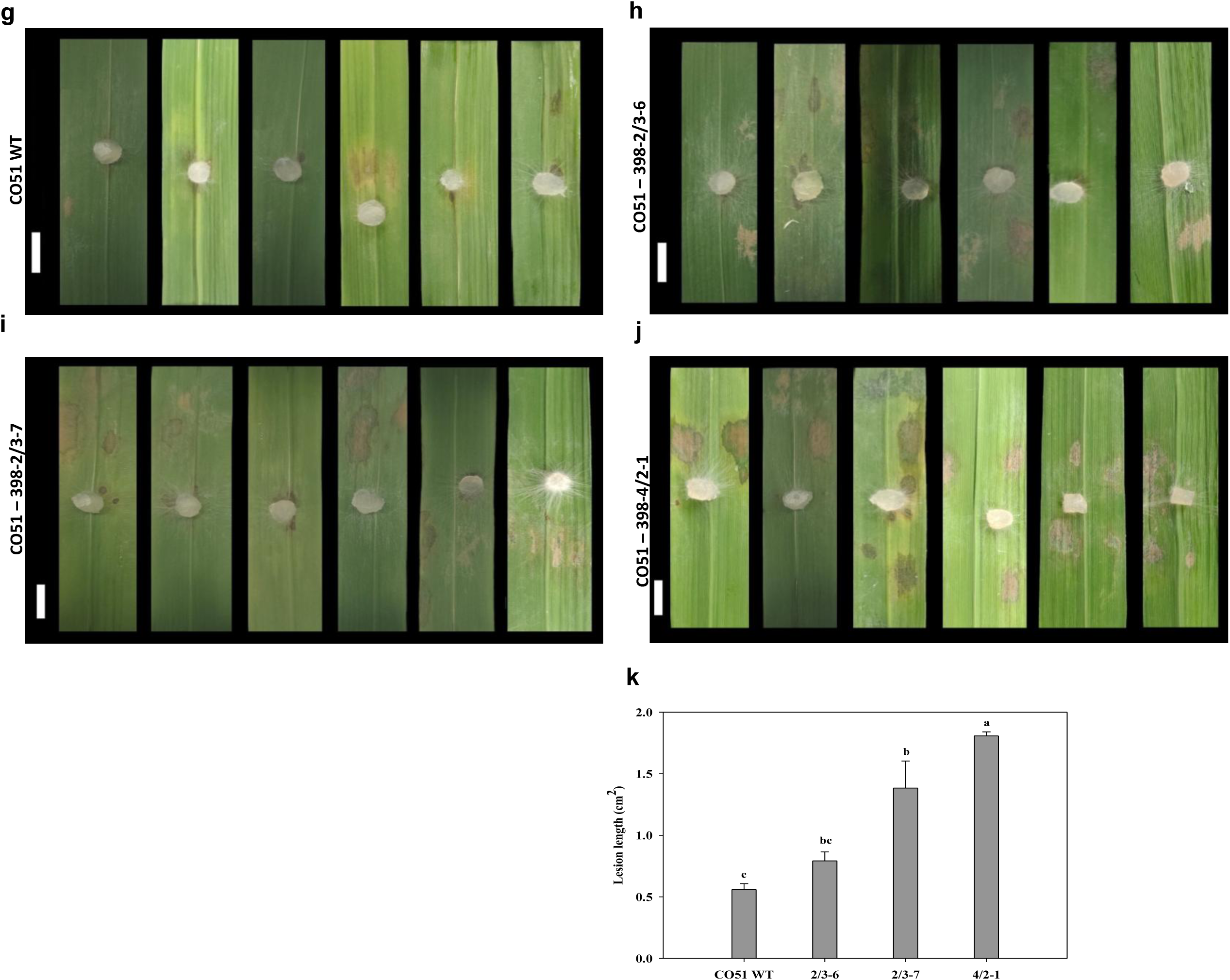
Assessment of detached leaf bioassay shows increased lesion formation in WT and *OsNH2* mutants after 48 hpi. (a-f) ASD16 and (g-k) CO51. Scale bar =1cm. Data represent means ± SEs, *n* = 10; Small letters above each bar show significant differences, detected by Tukey’s honestly significant difference test. The experiment was repeated three times with consistent results.

**Fig. S3.**
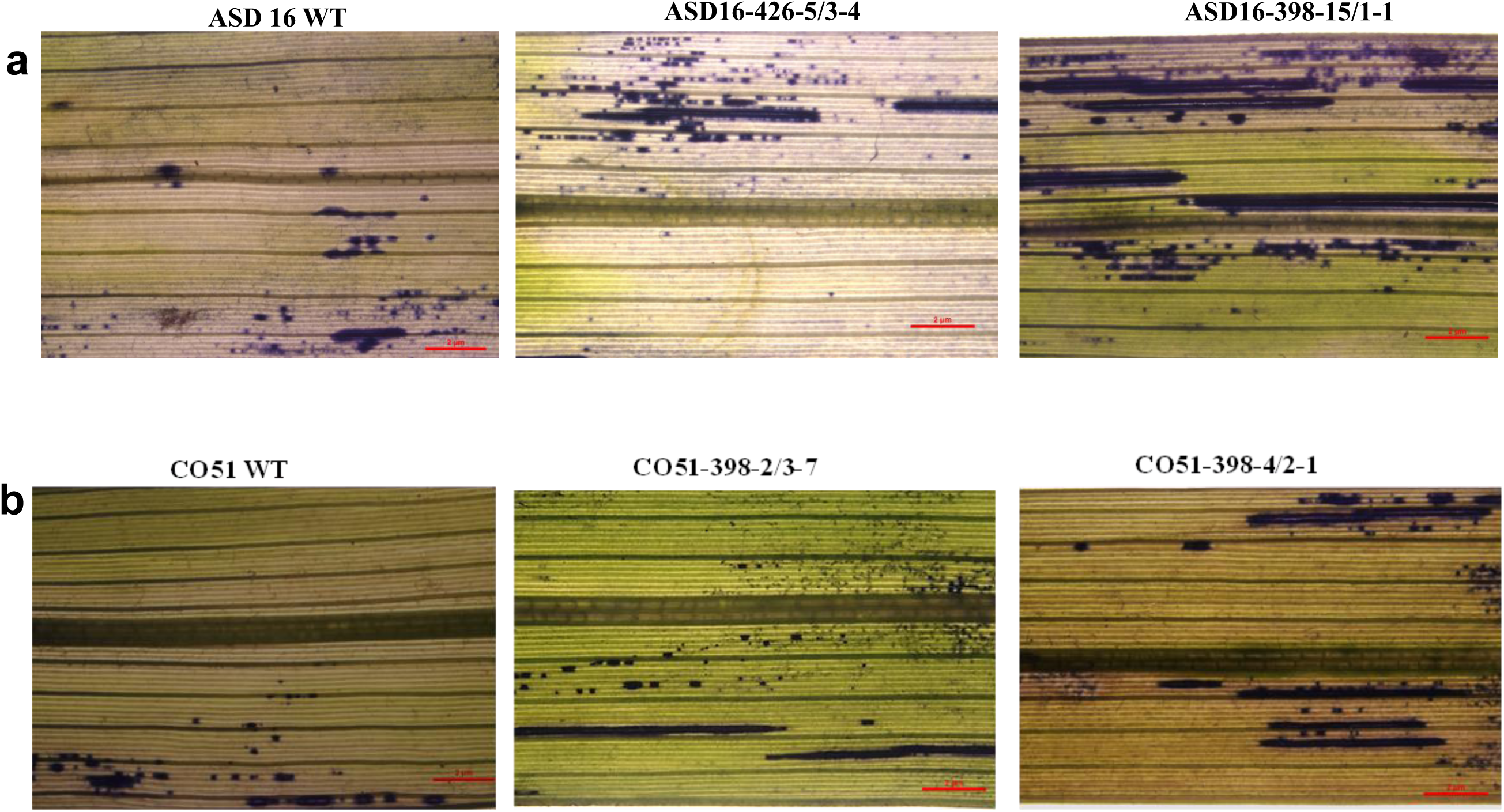
***In situ* detection of superoxide using NBT staining in WT plants and *OsNH2* mutants at 48 hpi**. (a) ASD16 and (b) CO51. Scale bar = 2 µm. The experiments were carried out in triplicate, showing consistent results.

**Fig. S4.**
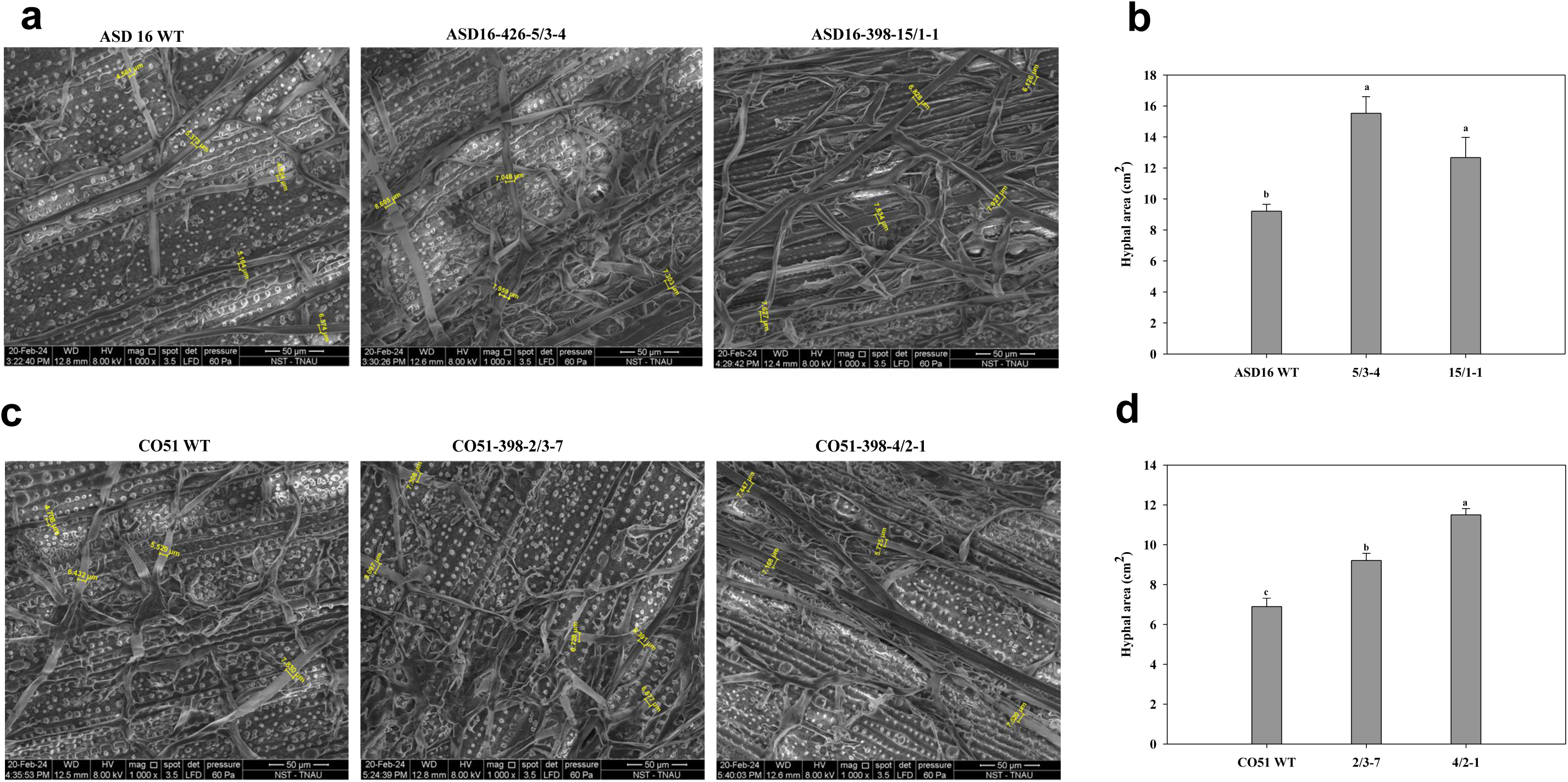
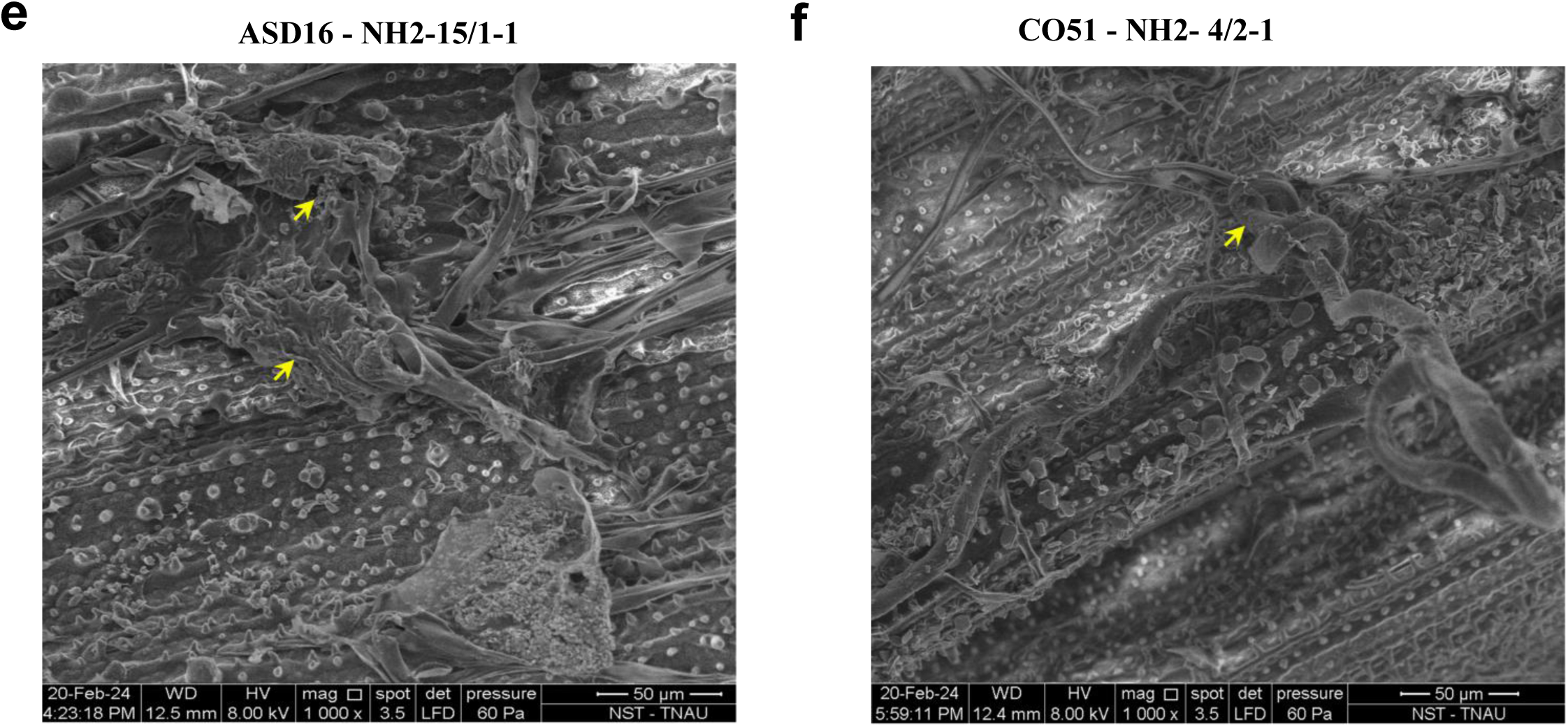
SEM images showing *R. solani* hyphal interactions on the leaf surfaces of WT plants and *OsNH2* mutants at 48 hpi. The bar graph illustrates the size of the hyphal area: (a, b) ASD16, (c, d) CO51, and (e, f) Infection cushions in *OsNH2* mutants. Scale bar = 50 µm. Data represent mean ± SE, *n* = 6. Lowercase letters above each bar show significant differences detected by Tukey’s honestly significant difference test. The hyphal area was measured using ImageJ software and plotted on a graph. Arrows indicate the infection cushions observed in mutant lines.

**Fig. S5.**
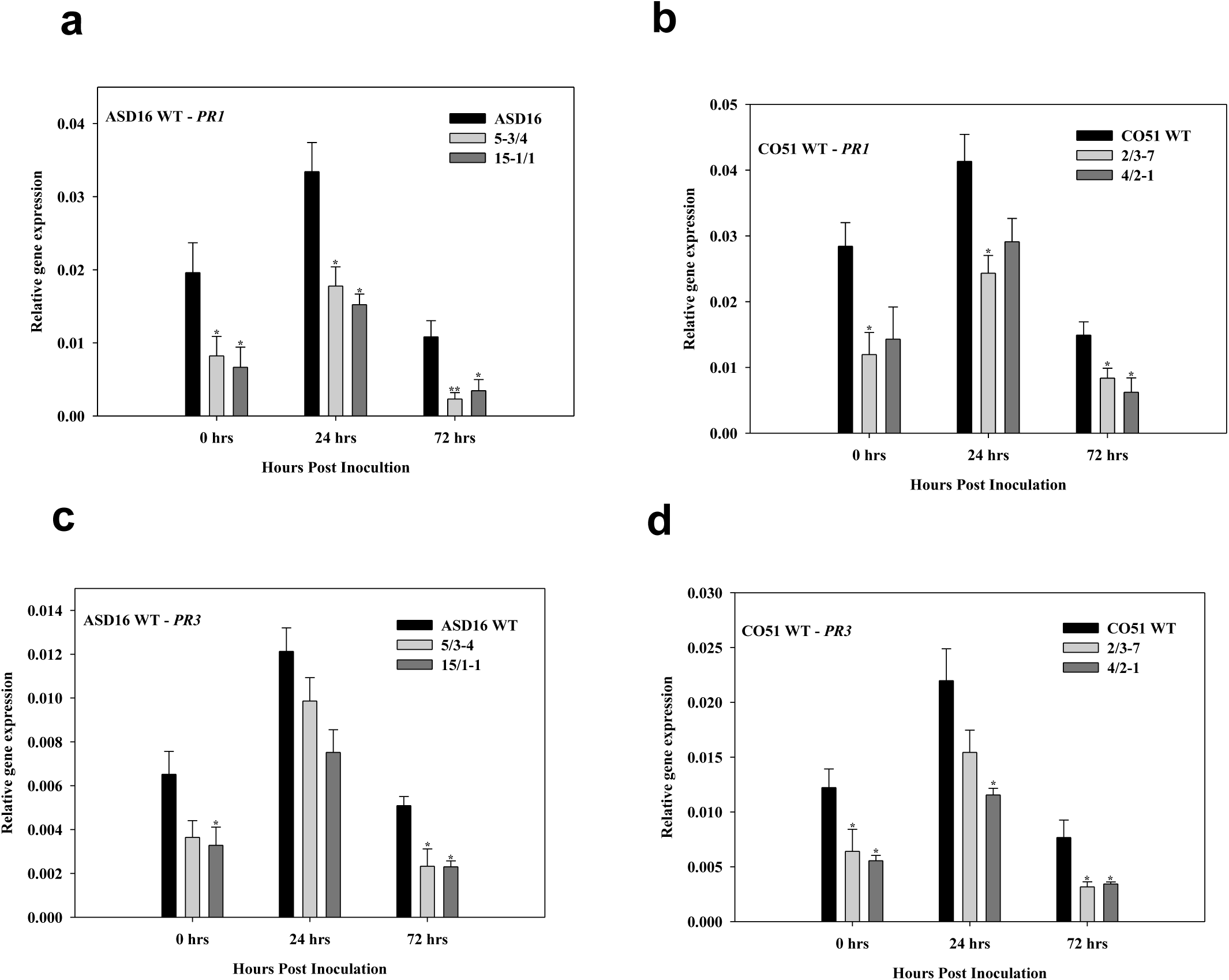
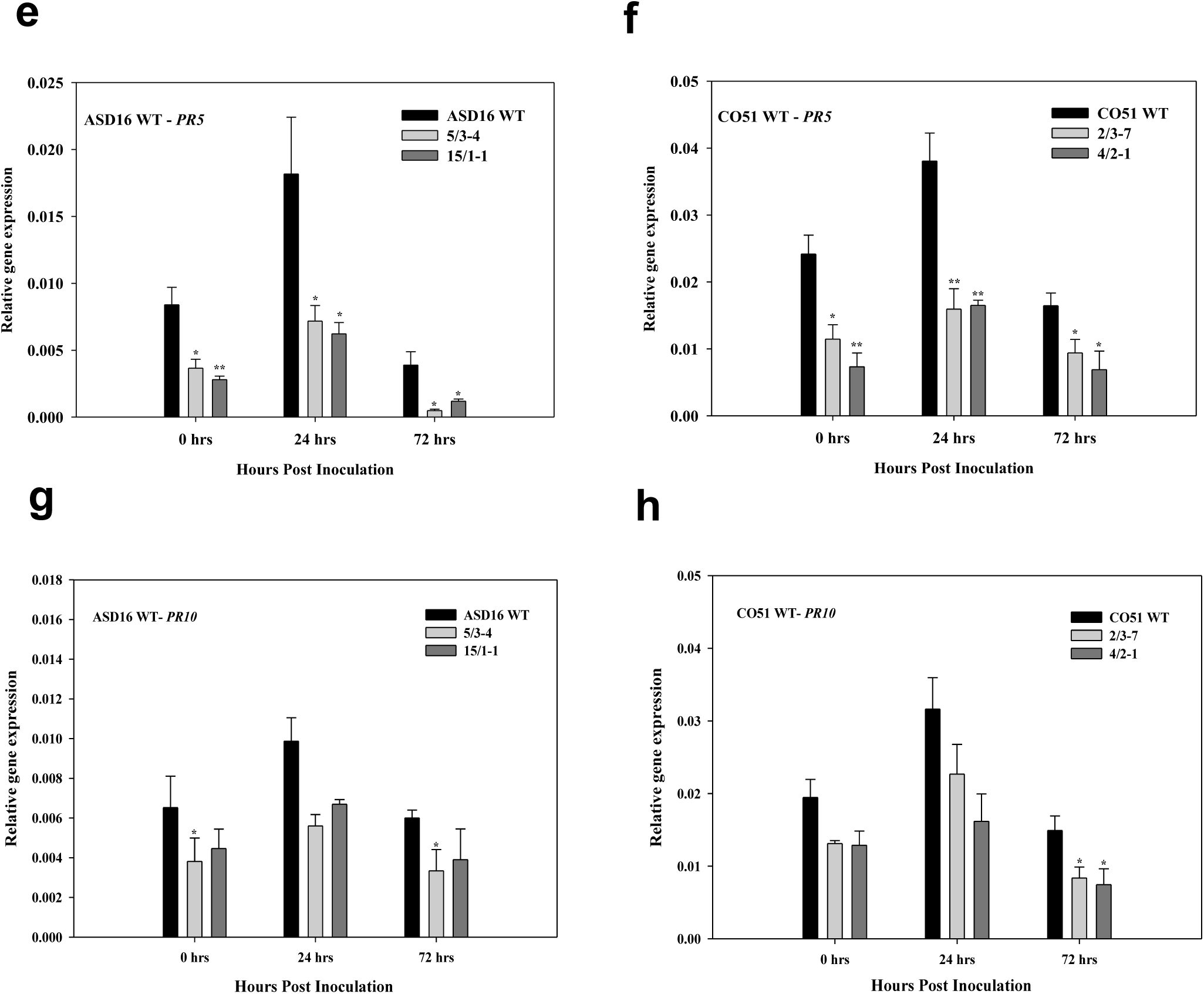
**Defense-related gene expression in WT plants and *OsNH2* mutants**. (a, b) PR1, (c, d) PR3, (e, f) PR5, and (g, h) PR10. Data represent mean ± SE. Asterisks indicate a significant difference (*P < 0.05, **P < 0.01, ***P < 0.001 Student’s t-test) between different time intervals. The experiment was carried out in duplicate with consistent results.

**Fig. S6.**
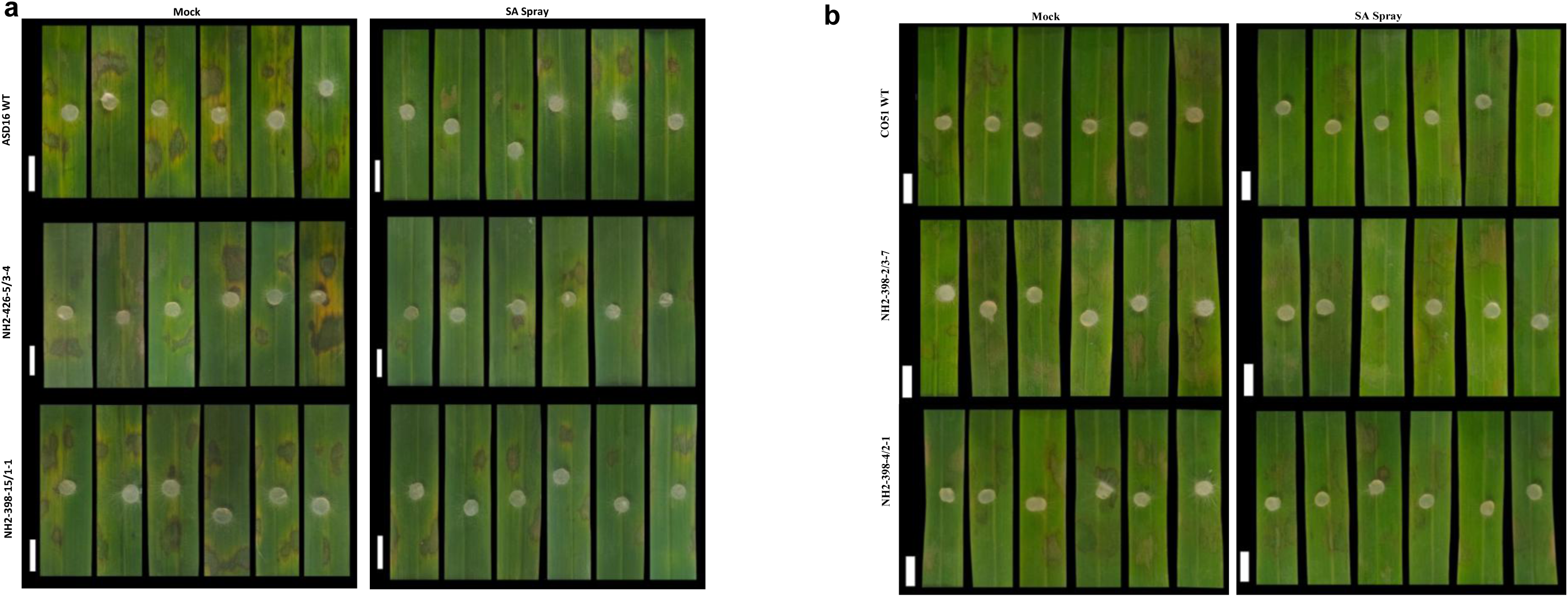
**Comparison of lesion formation after exogenous SA application and mock treatment in WT plants and their mutants. (**a) ASD16, and (b) CO51 in detached leaf assay after 48 hpi.

**Fig. S7.**
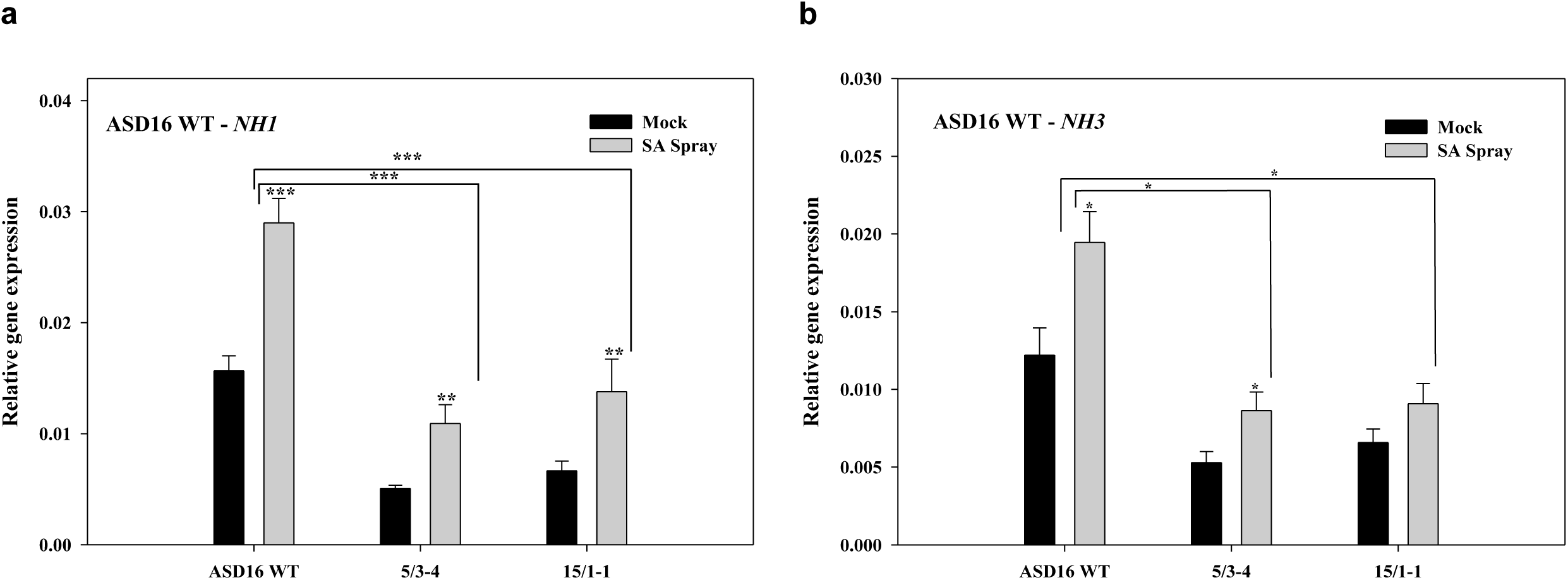
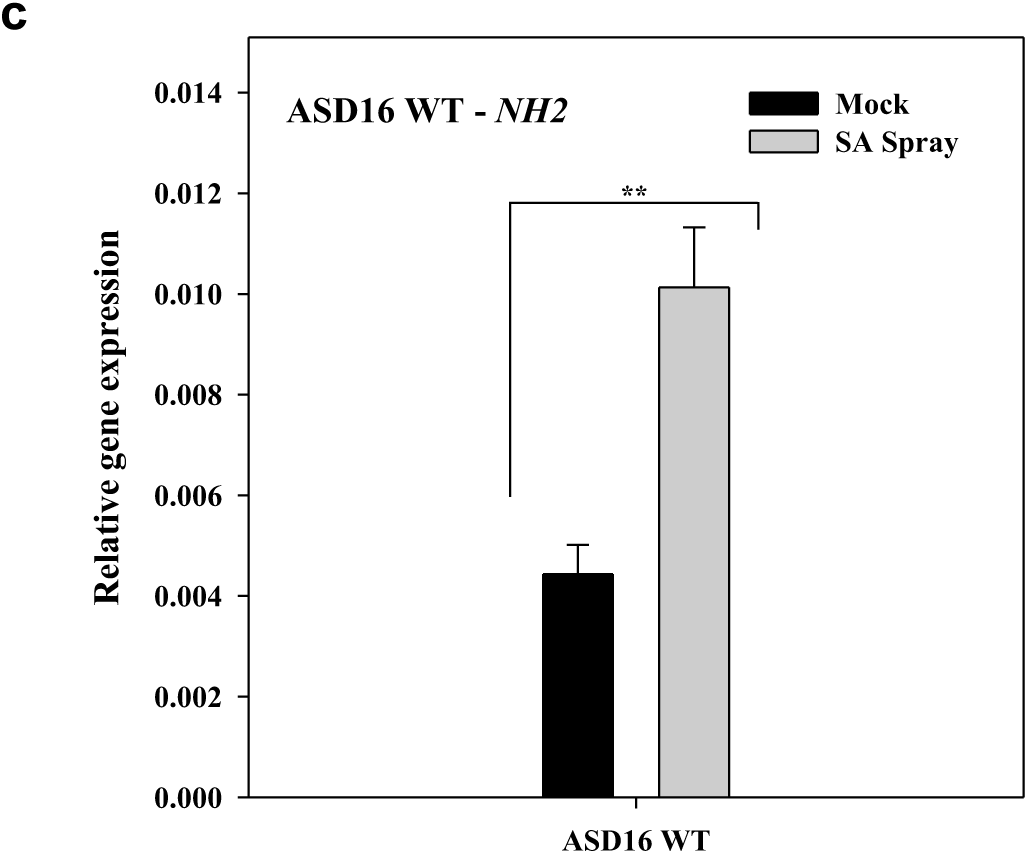
Expression of *OsNH1* and *OsNH3* in ASD16 WT plants and *OsNH2* mutants after SA spray. Expression of (a) *OsNH1* and (b) *OsNH3* was highly induced 24 h after SA spray, followed by *R. solani* inoculation, (c) Strong induction of *OsNH2* expression in ASD16 WT plants after SA treatment at 24 hpi. Data represent mean ± SE. Asterisks indicate a significant difference (*P < 0.05, **P< 0.01, ***P< 0.001 Student’s t-test). The experiment was carried out in triplicate with similar trends among each other.

**Fig. S8.**
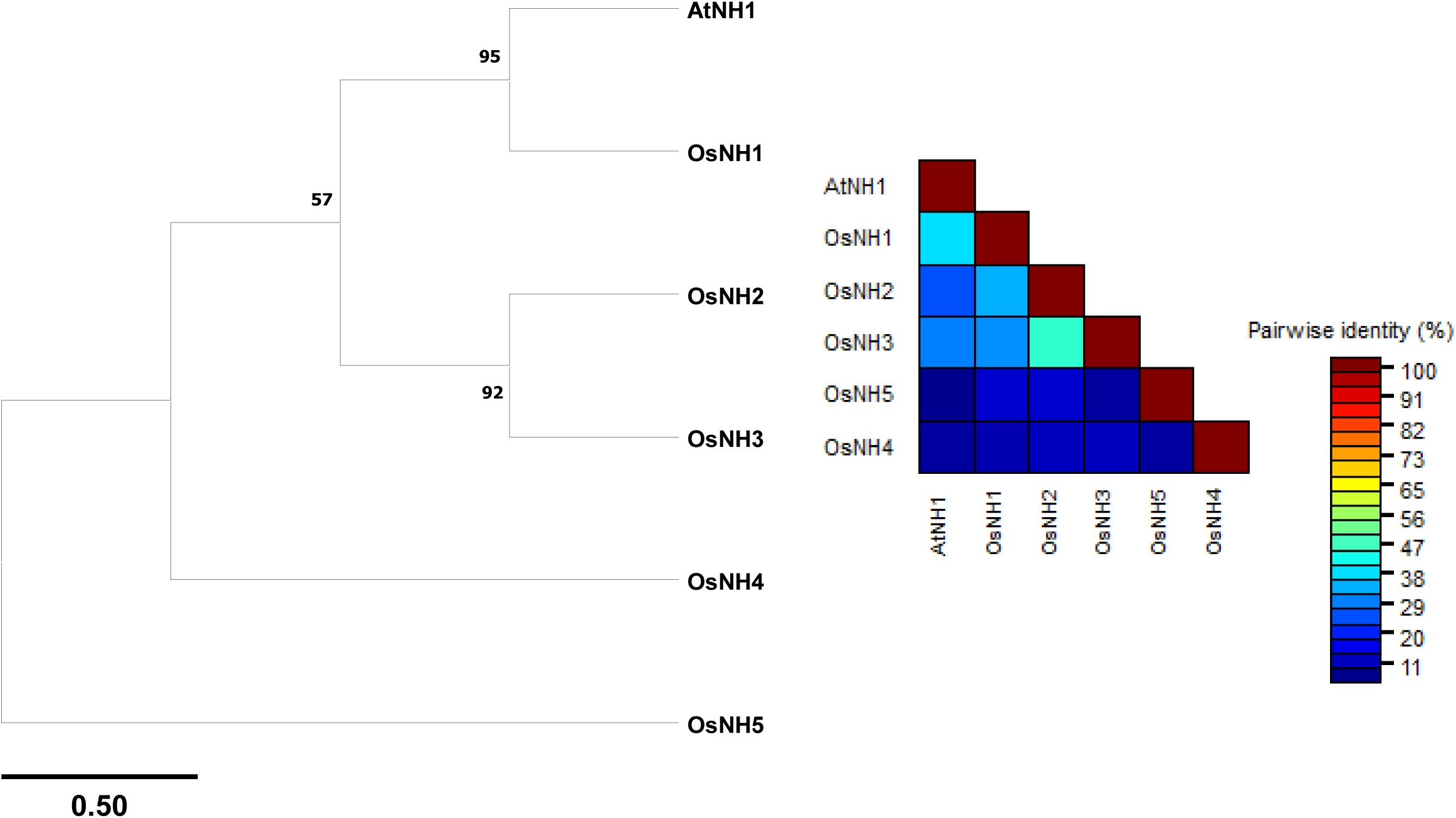
Phylogenetic tree of NPR1-like proteins in rice and Arabidopsis. The phylogenetic tree was constructed using the sequences of five rice OsNH gene family proteins (*OsNH1* to *OsNH5*) and AtNH1 from Arabidopsis. Protein sequences were aligned using BlastP, and the percentage of amino acid (aa) identity was calculated by comparing each rice protein to AtNH1/NPR1. This analysis highlights the evolutionary relationships and sequence conservation among NPR1-like proteins.

**Fig. S9.**
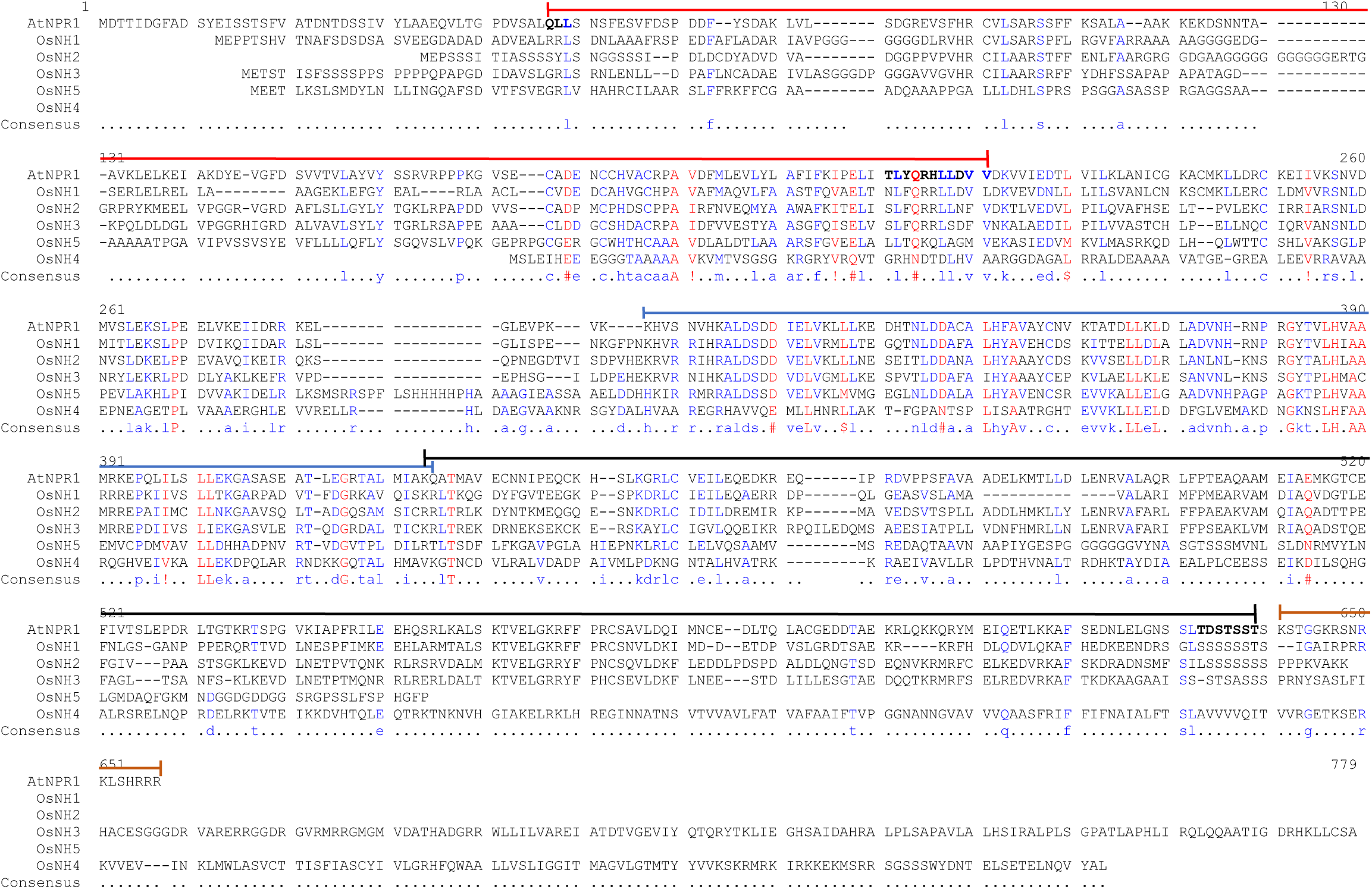
Multiple sequence alignment of the five OsNH proteins (*OsNH1, OsNH2, OsNH3, OsNH4, and OsNH5*) alongside *AtNPR1*, highlighting key functional domains critical for their roles in plant immune signaling. The alignment is color-coded to highlight specific domains that contribute to the structure and function of these proteins: **Red** indicates the BTB/POZ domain (amino acids 47–194) for protein interactions and signal transduction, **Blue** represents the ankyrin repeats (amino acids 265–371) for transcription factor interactions, **Black** denotes the NPR1/NIM1-like defense protein C-terminal region (amino acids 370–575) for defense gene activation, and **Brown** highlights the nuclear localization signal (amino acids 577–593) for nuclear targeting and immune response activation. This alignment underscores the conservation of these domains across the OsNH proteins and *AtNH1*, reflecting their evolutionary significance and functional importance in SA-mediated defense pathways.

**Fig. S10.**
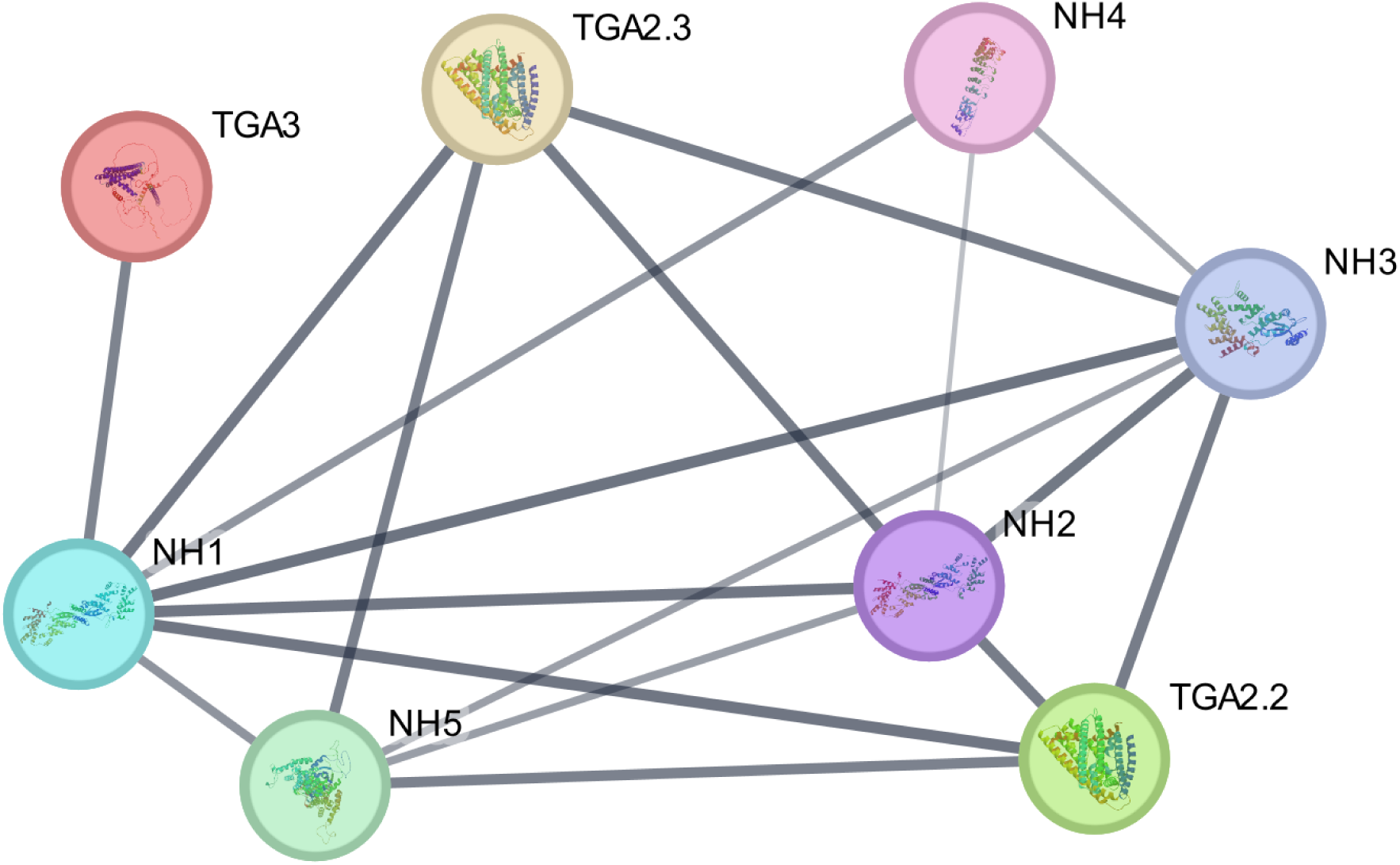
STRING-based protein–protein interaction network of NH gene family members and TGA transcription factors in rice. The network illustrates both predicted and experimentally supported interactions among NH1 to NH5 and the TGA transcription factors TGA2.2, TGA2.3, and TGA3. A highly interconnected subnetwork is formed by *NH1*, *NH2*, and *NH3*, which show strong associations with multiple TGA factors, suggesting potential co-regulatory or functional interactions in rice immunity. The thickness of each edge reflects the strength of supporting evidence, with thicker edges indicating higher STRING confidence scores.

**Fig. S11.**
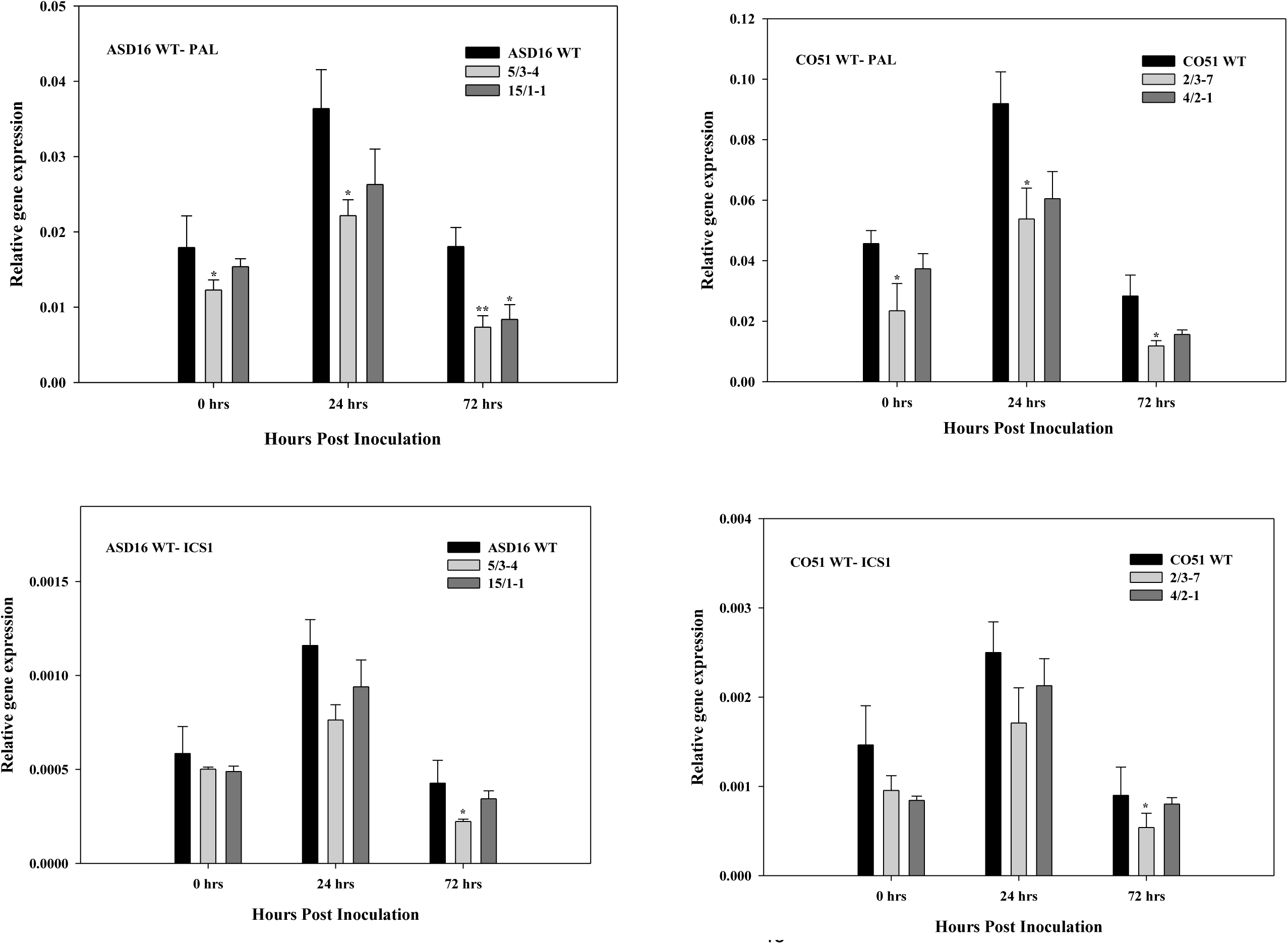
Expression of key SA biosynthesis genes in WT and *OsNH2* mutants. (a, b) Expression of PAL, and (c, d) ICS1 at different time points. Data are presented as mean ± SE. Asterisks denote statistically significant differences compared to corresponding time points (*P* < 0.05, P < 0.01, Student’s *t*-test). The experiment was independently repeated twice with similar results.

**Fig. S12.**
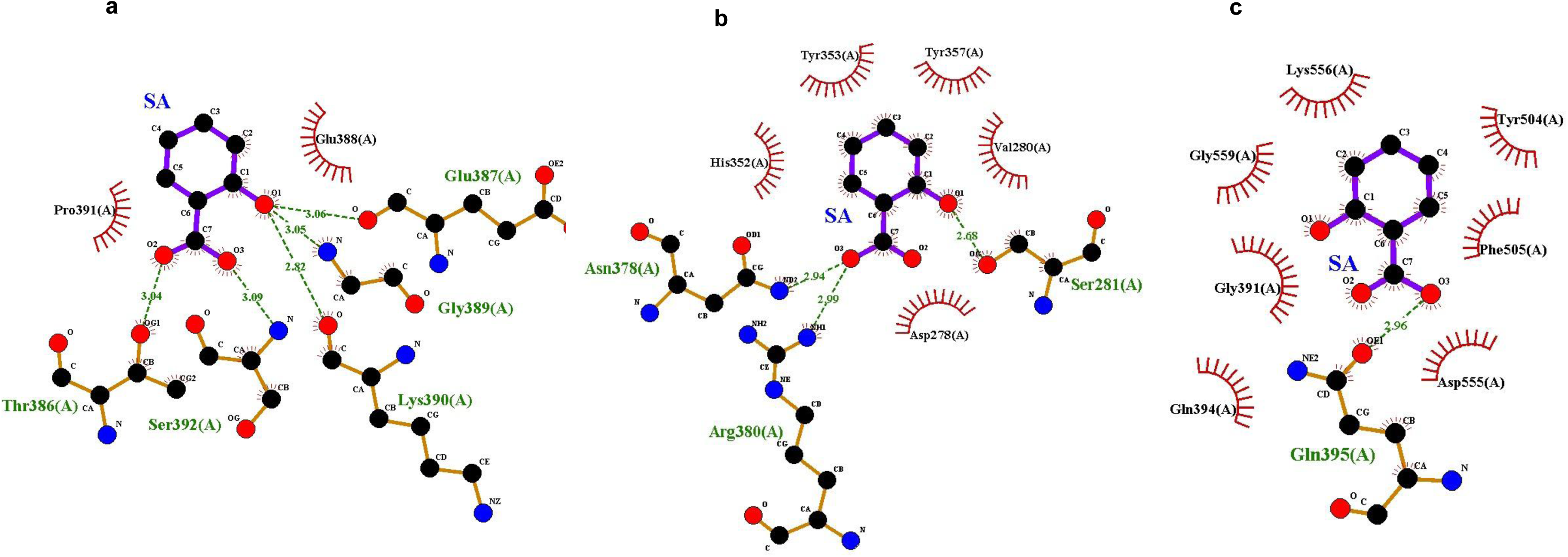
Predicted interaction of SA with SA-binding domains of NPR1 homologs OsNH1, OsNH2, and OsNH3. Molecular docking analysis shows that SA binds within the C-terminal domain of (a) OsNH1 (amino acids 391–526), (b) OsNH2 (439–576), and (c) OsNH3 (382–521). Green indicates amino acid residues involved in hydrogen bonding, with dotted lines representing hydrogen bonds and their lengths shown in angstroms (Å). Brown highlights residues involved in van der Waals interactions.

**Table S1.**
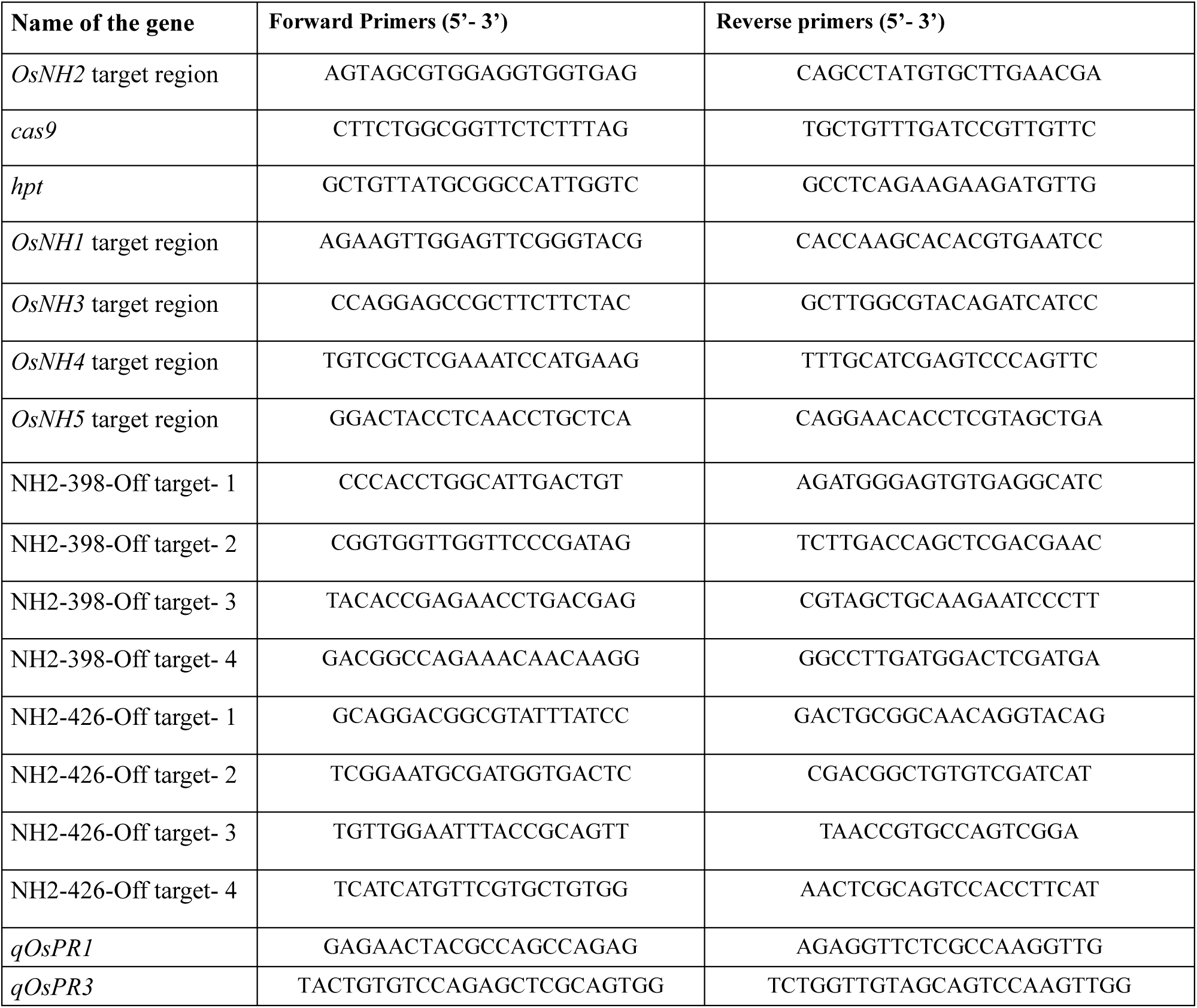
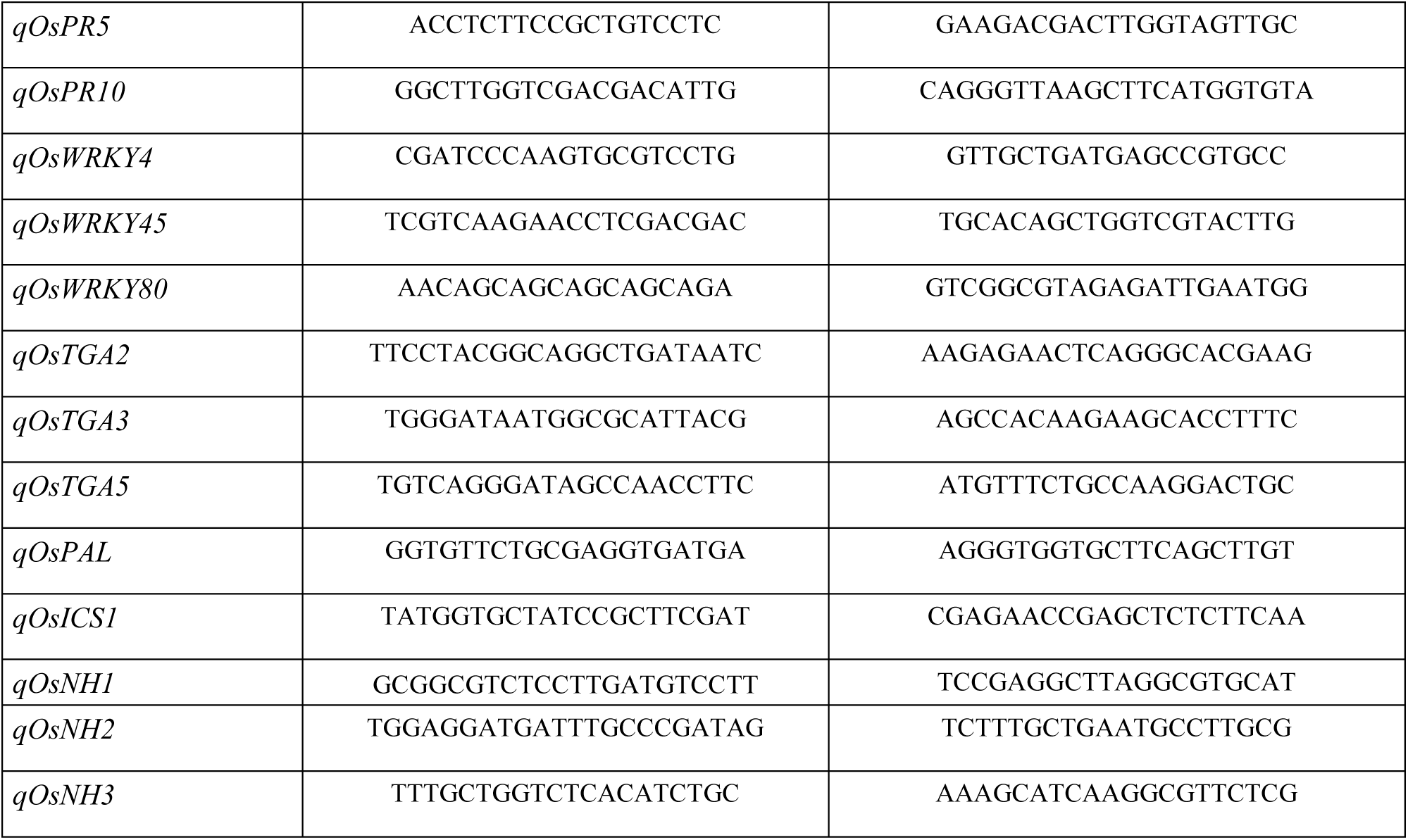
List of primers used in this study

**Table S2.**
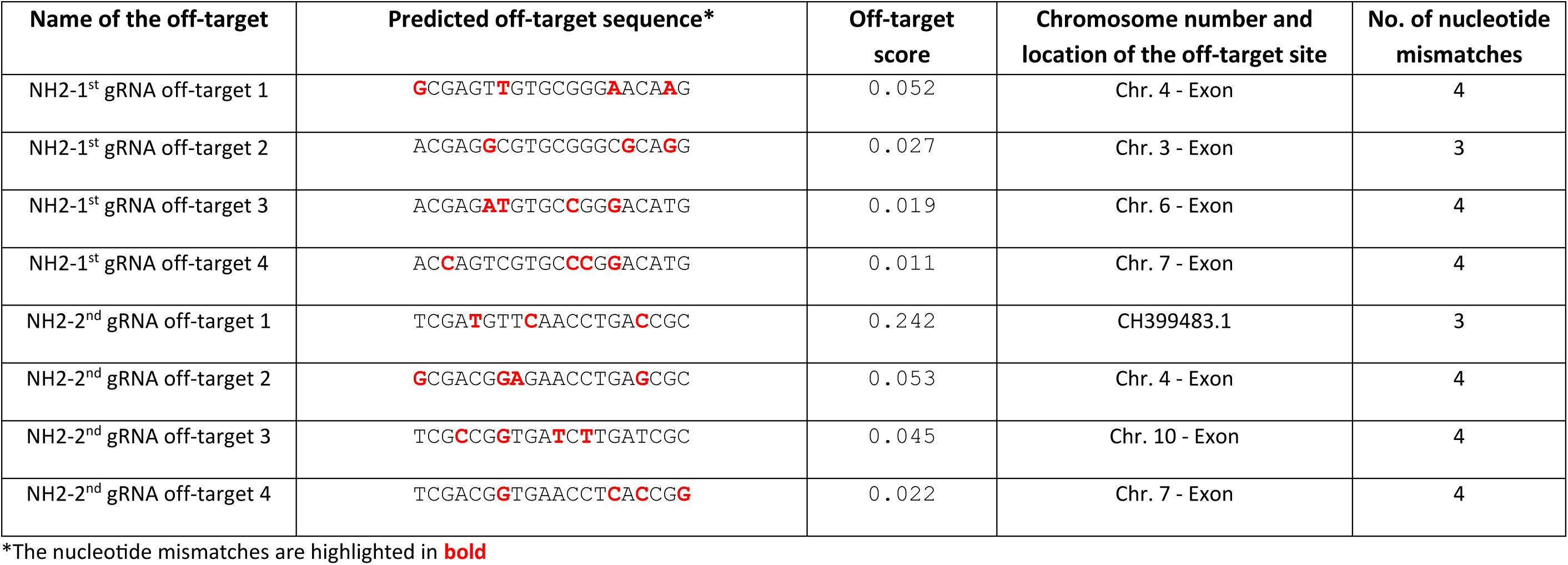
List of predicted off-target sites

**Table S3.** Transformation and mutation efficiency of *Agrobacterium*-mediated genetic transformation of rice cultivars ASD16 and CO51 using CRISPR-*OsNH2* constructs.

| Genotype | Construct Name | Number of embryos co-cultivated | Number of events generated | Number of events with mutation | Transformation efficiency (%) | Mutation efficiency (%) |
| --- | --- | --- | --- | --- | --- | --- |
| ASD16 | NH2 -398 | 1020 | 20 | 12 | 1.96 | 60.00 |
|  | NH2 -426 | 1500 | 40 | 17 | 2.67 | 42.50 |
| CO51 | NH2 -398 | 600 | 15 | 10 | 2.50 | 66.67 |
| Total |  |  |  |  |  | 56.39 |

The efficiency of *Agrobacterium*-mediated transformation was calculated using the formula,
Based on the sequencing result analysis, mutation efficiency was calculated as,

**Table S4.**
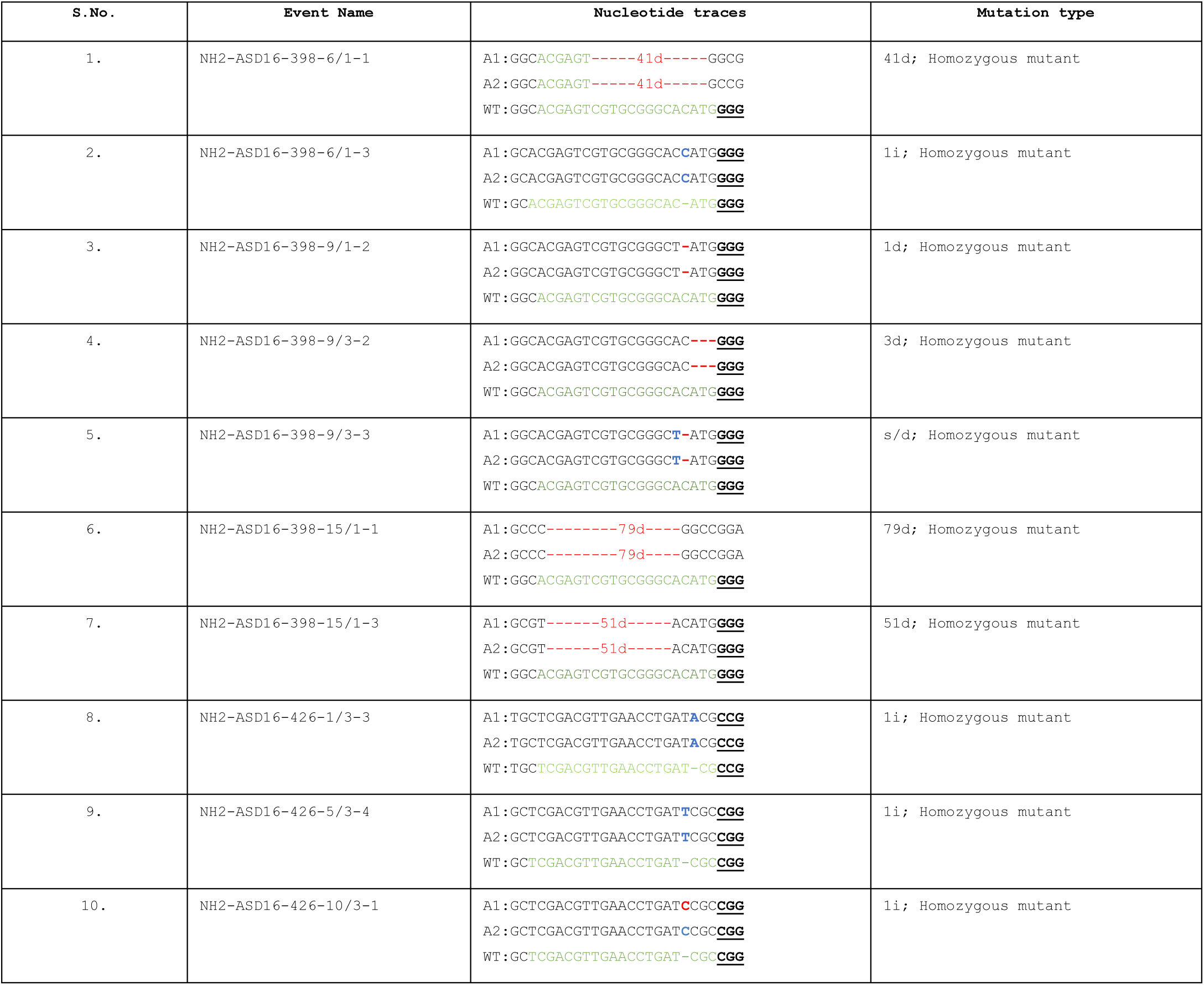
Mutation and zygosity of *OsNH2* mutants observed in T_1_ generation (ASD16)

**Table S5.**
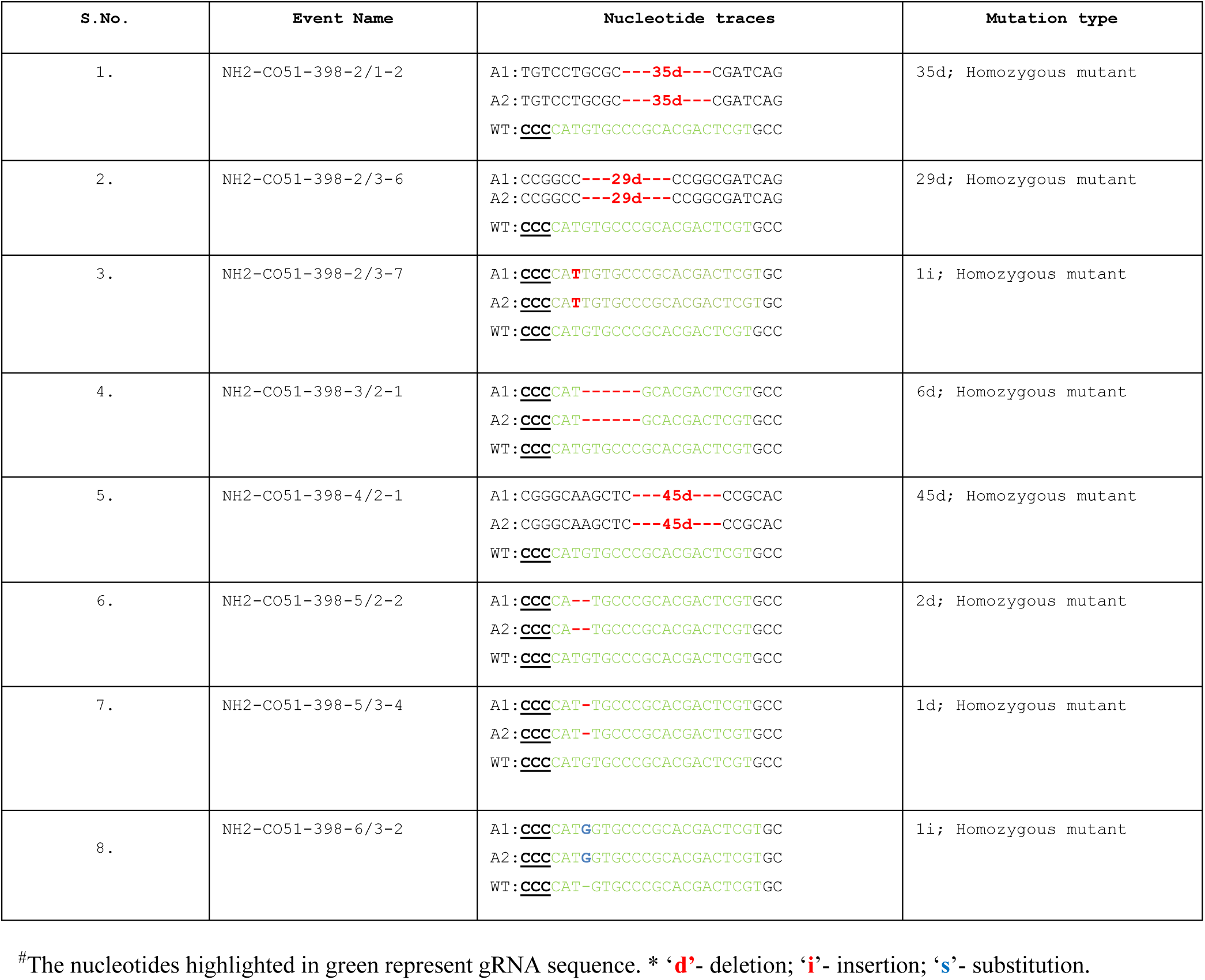
Mutation and zygosity of *OsNH2* mutants observed in T_1_ generation (CO51)

**Table S6.**
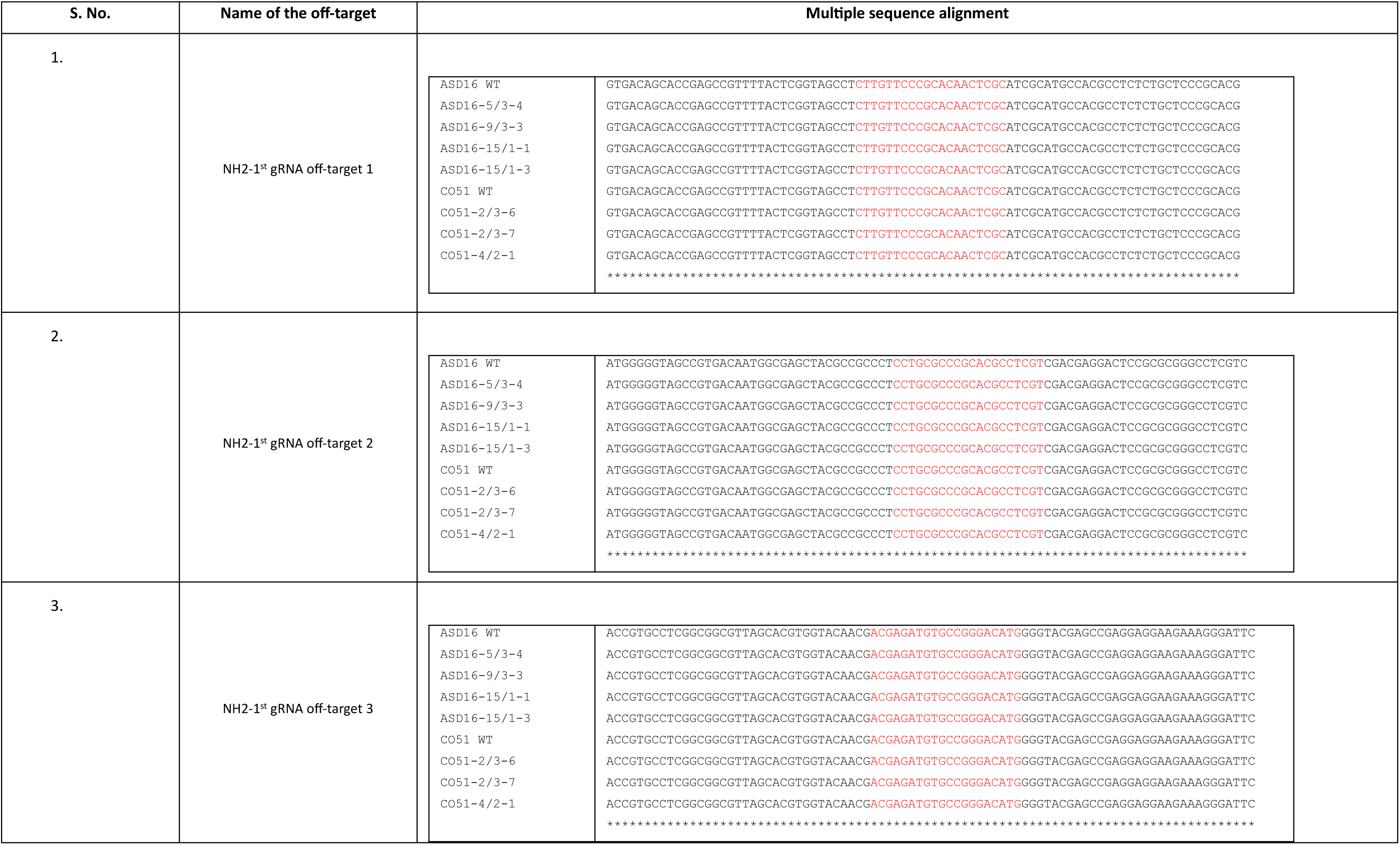
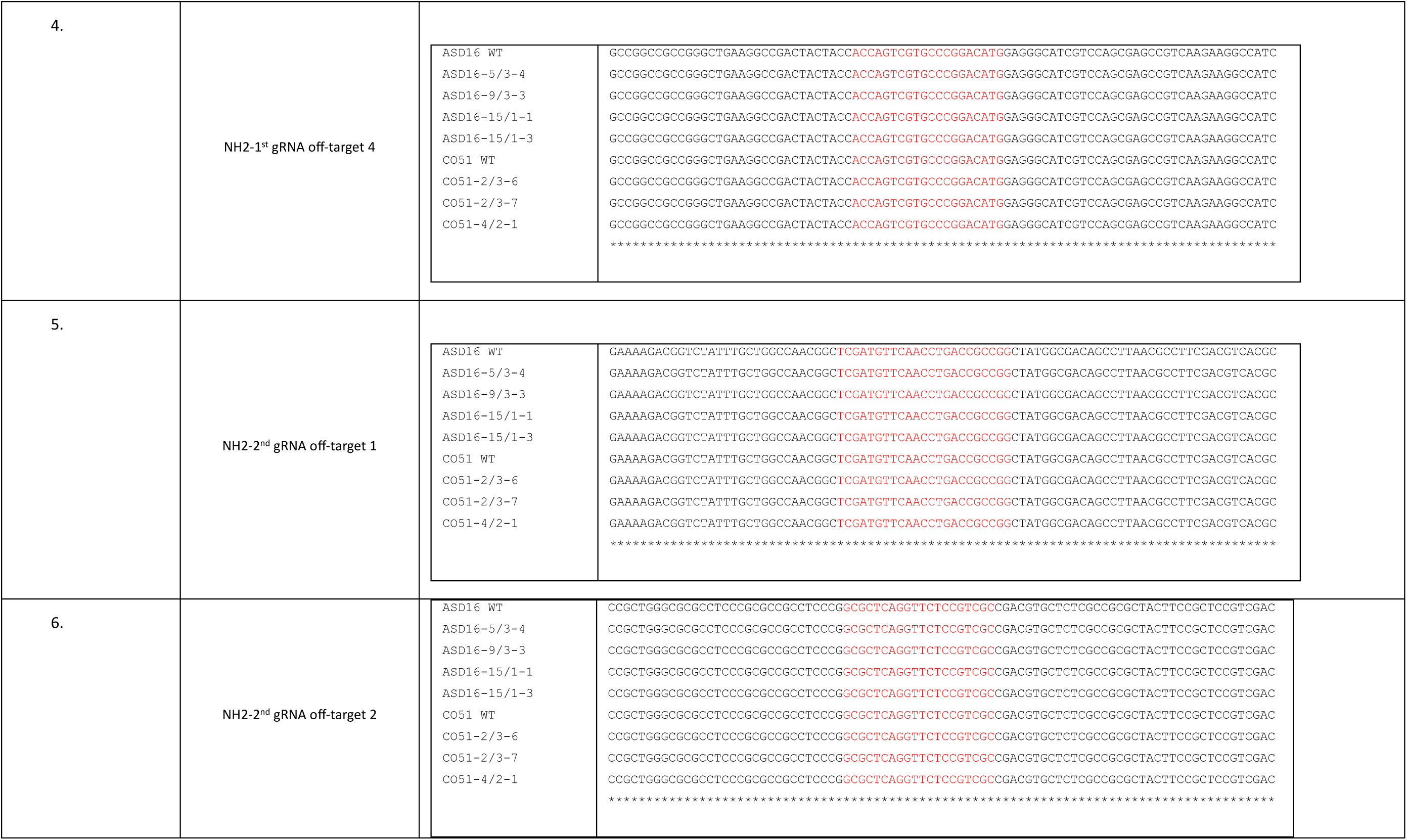
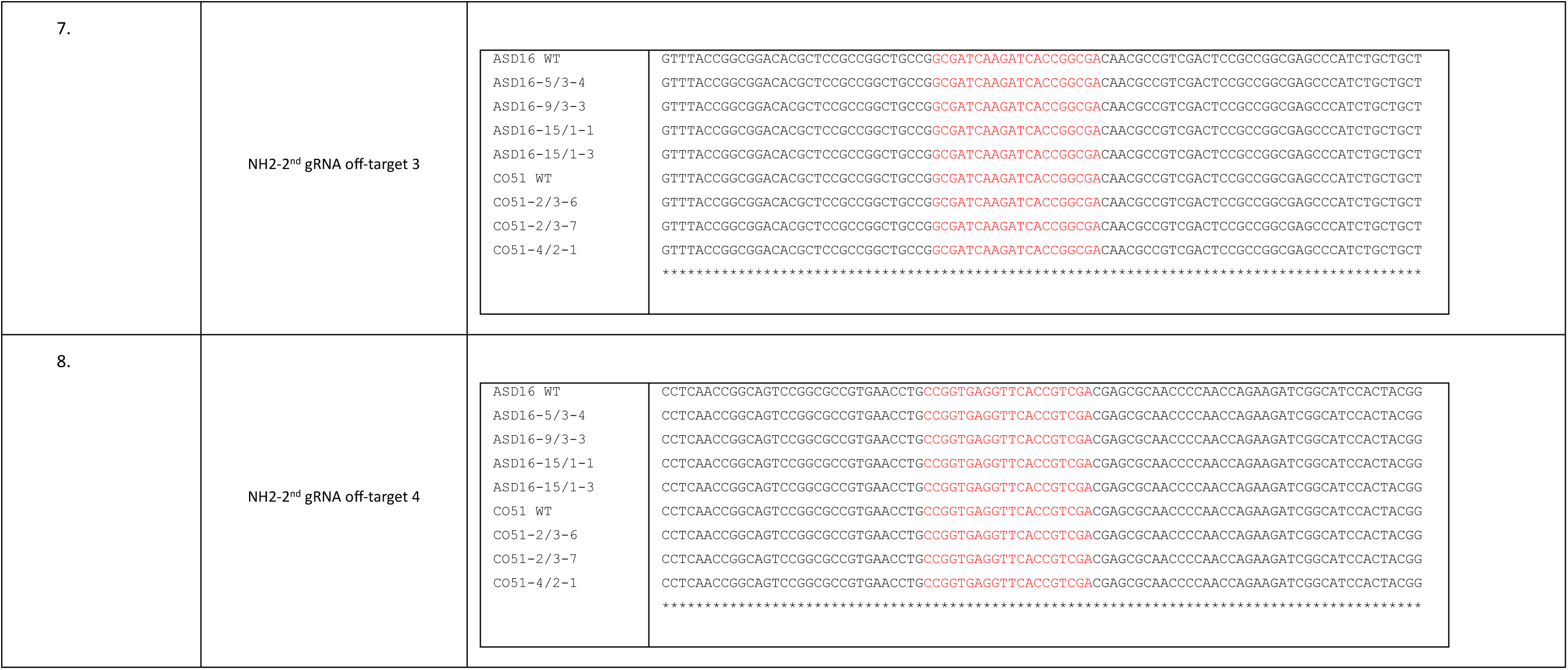
Off-target analysis of *OsNH2* mutants in ASD16 and CO51 cultivars

**Table S7.**
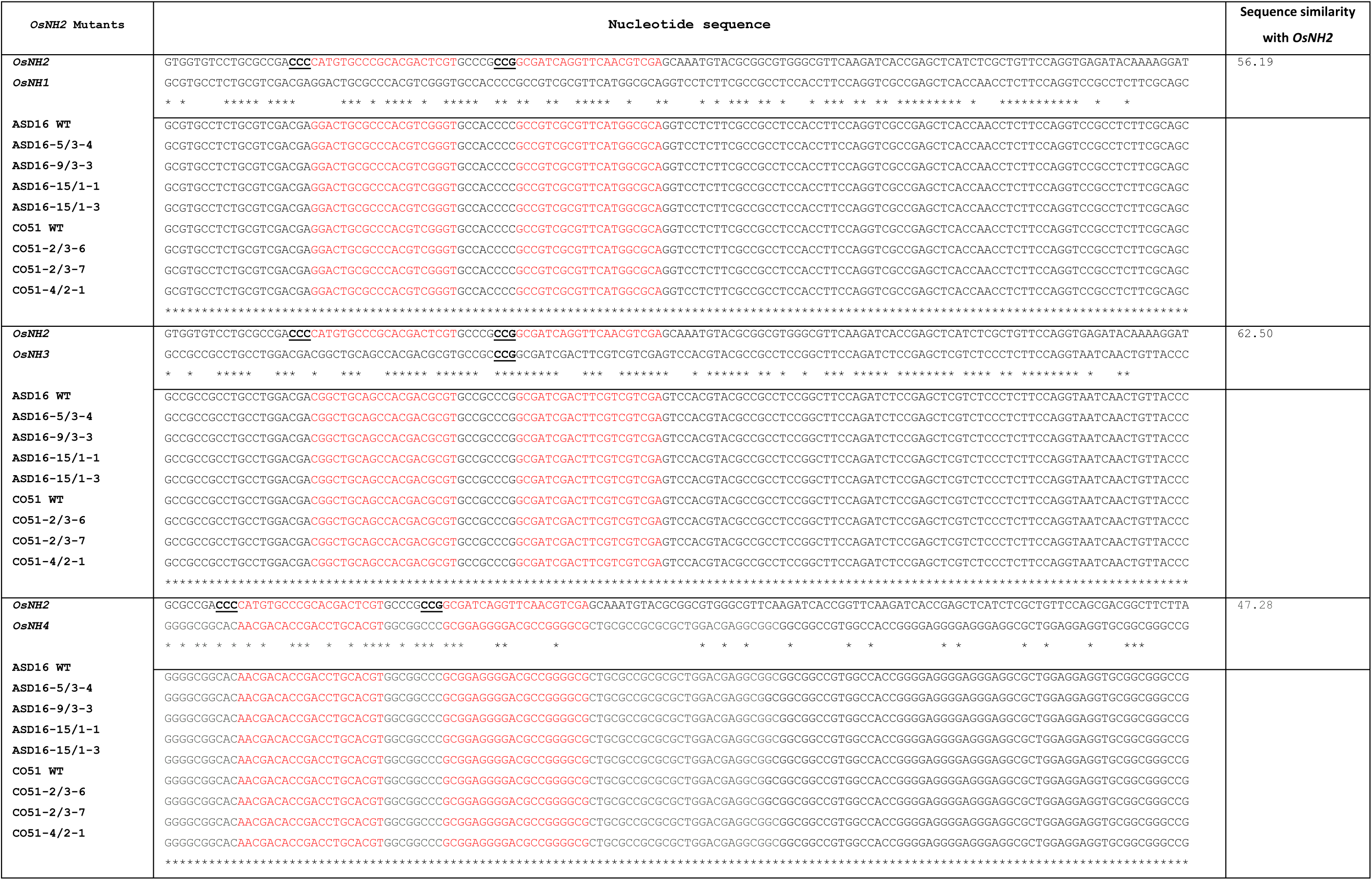
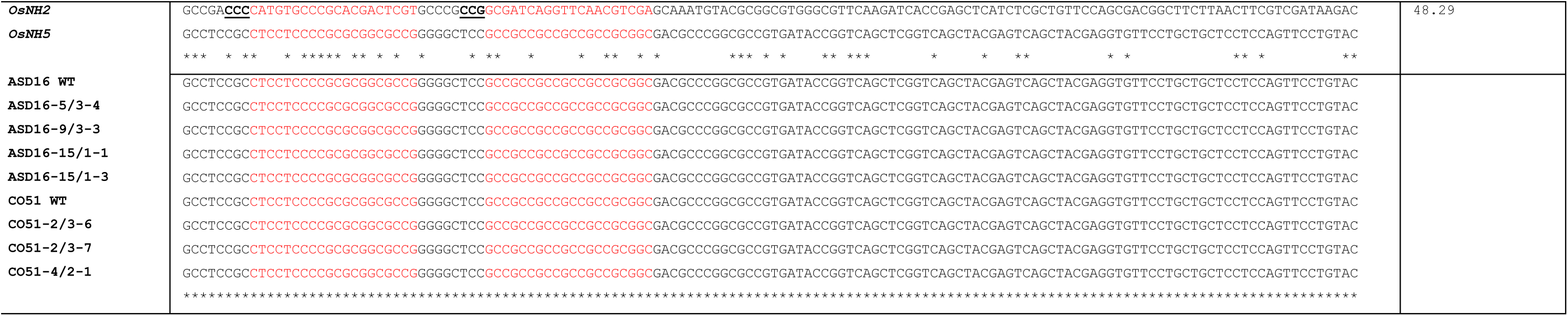
Off-target analysis of *OsNH1*, *OsNH3*, *OsNH4*, and *OsNH5* in *OsNH2* mutants. MSA revealed sequence similarity among *OsNH* gene family members with *OsNH2*

**Table S8.**
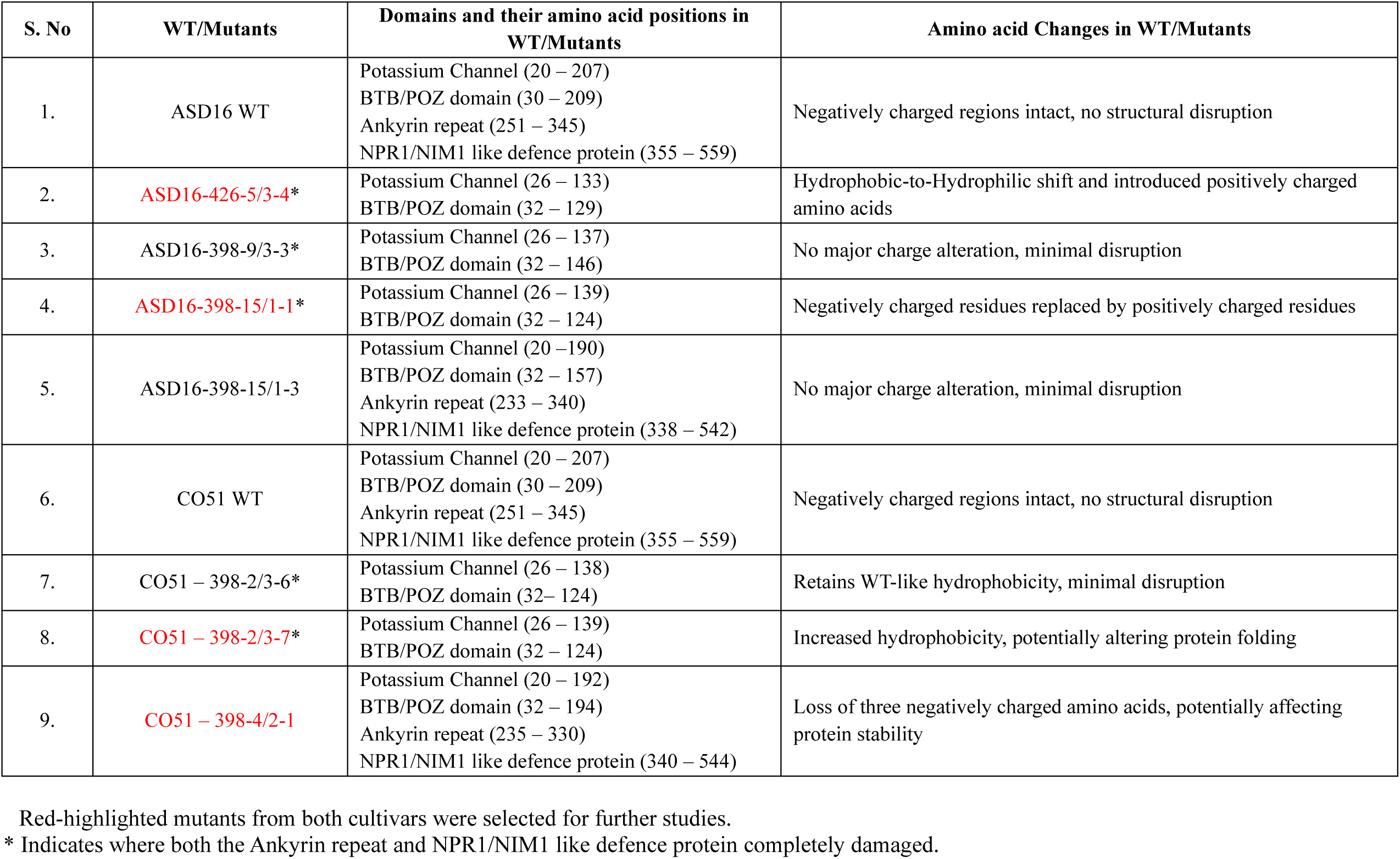
List of domain disruptions and amino acid changes in WT and *OsNH2* mutants

## Notes

### Competing Interest Statement

The authors have declared no competing interest.

## References

1. Aravind L, Koonin EV (1999) Fold prediction and evolutionary analysis of the POZ domain: structural and evolutionary relationship with the potassium channel tetramerization domain. J. Mol. Biol. 285(4):1353–1361. 10.1006/jmbi.1998.2394

2. Backer R, Naidoo S, Van den Berg N (2019) The NONEXPRESSOR OF PATHOGENESIS-RELATED GENES 1 (NPR1) and related family: mechanistic insights in plant disease resistance. Front. Plant. Sci. 10:102. 10.3389/fpls.2019.00102

3. Bai W, Chern M, Ruan D, Canlas PE, Sze-to WH, Ronald PC (2011) Enhanced disease resistance and hypersensitivity to BTH by introduction of an NH1/*OsNPR1* paralog. Plant Biotechnol. J. 9(2):205–215. 10.1111/j.1467-7652.2010.00544.x

4. Bari R, Jones JD (2009) Role of plant hormones in plant defence responses. Plant Mol. Biol. 69:473–488. 10.1007/s11103-008-9435-0

5. Basu A, Chowdhury S, Ray Chaudhuri T, Kundu S (2016) Differential behaviour of sheath blight pathogen *Rhizoctonia solani* in tolerant and susceptible rice varieties before and during infection. Plant Pathol. 65(8):1333–1346. 10.1111/ppa.12502

6. Boyle P, Le Su E, Rochon A, Shearer HL, Murmu J, Chu JY, Fobert PR, Després C (2009) The BTB/POZ domain of the Arabidopsis disease resistance protein NPR1 interacts with the repression domain of TGA2 to negate its function. Plant Cell 21(11):3700–3713. 10.1105/tpc.109.069971

7. Cao H, Bowling SA, Gordon AS, Dong X (1994) Characterization of an *Arabidopsis* mutant that is nonresponsive to inducers of systemic acquired resistance. Plant Cell 6(11):1583–1592. 10.1105/tpc.6.11.1583

8. Cao H, Glazebrook J, Clarke JD, Volko S, Dong X (1997) The *Arabidopsis* NPR1 gene that controls systemic acquired resistance encodes a novel protein containing ankyrin repeats. Cell 88(1):57–63. 10.1016/S0092-8674(00)81858-9

9. Cao H, Li X, Dong X (1998) Generation of broad-spectrum disease resistance by overexpression of an essential regulatory gene in systemic acquired resistance. Proc. Natl. Acad. Sci. USA. 95:6531–6536. 10.1073/pnas.95.11.6531

10. Cao W, Zhang H, Zhou Y, Zhao J, Lu S, Wang X, Chen X, Yuan L, Guan H, Wang G, Shen W (2022) Suppressing chlorophyll degradation by silencing *OsNYC3* improves rice resistance to *Rhizoctonia solani*, the causal agent of sheath blight. Plant Biotechnol. J. 20(2):335–349. 10.1111/pbi.13715

11. Chen H, Lin Q, Li Z, Chu J, Dong H, Mei Q, Xuan Y (2023) Calcineurin B-like interacting protein kinase 31 confers resistance to sheath blight via modulation of ROS homeostasis in rice. Mol. Plant. Pathol. 24(3):221–231. 10.1111/mpp.13291

12. Chen J, Clinton M, Qi G, Wang D, Liu F, Fu ZQ (2020) Reprogramming and remodeling: transcriptional and epigenetic regulation of salicylic acid-mediated plant defense. J. Exp. Bot. 71(17):5256–5268. 10.1093/jxb/eraa072

13. Chern M, Bai W, Ruan D, Oh T, Chen X, Ronald PC (2014) Interaction specificity and coexpression of rice NPR1 homologs 1 and 3 (NH1 and NH3), TGA transcription factors and Negative Regulator of Resistance (NRR) proteins. BMC genomics 15:1–20. 10.1186/1471-2164-15-461

14. Chern M, Fitzgerald HA, Canlas PE, Navarre DA, Ronald PC (2005) Overexpression of a rice NPR1 homolog leads to constitutive activation of defense response and hypersensitivity to light. Mol. Plant. Microbe Interact. 18(6):511–520. 10.1094/MPMI-18-0511

15. Cui Z, Xue C, Mei Q, Xuan Y (2022) Malectin Domain Protein Kinase (MDPK) Promotes Rice Resistance to Sheath Blight via IDD12, IDD13, and IDD14. Int. J. Mol. Sci. 23(15):8214. 10.3390/ijms23158214

16. Dai X, Wang Y, Yu K, Zhao Y, Xiong L, Wang R, Li S (2023) *OsNPR1* enhances rice resistance to *Xanthomonas oryzae* pv*. oryzae* by upregulating rice defense genes and repressing bacteria virulence genes. Int. J. Mol. Sci. 24(10):8687. 10.3390/ijms24108687

17. Dehairs J, Talebi A, Cherif Y, Swinnen JV (2016) CRISP-ID: Decoding CRISPR mediated indels by Sanger sequencing. Sci. Rep. 6(1):28973. 10.1038/srep28973

18. Ding Y, Sun T, Ao K, Peng Y, Zhang Y, Li X, Zhang Y (2018) Opposite roles of salicylic acid receptors NPR1 and NPR3/NPR4 in transcriptional regulation of plant immunity. Cell 173(6):1454–1467. 10.1016/j.cell.2018.03.044

19. Farooq M, Basra SMA, Wahid A, Ahmad N, Saleem BA (2009) Improving the drought tolerance in rice (*Oryza sativa* L.) by exogenous application of salicylic acid. J. Agron. Crop. Sci. 195(4):237–246. 10.1111/j.1439-037X.2009.00365.x

20. Fister AS, Landherr L, Maximova SN, Guiltinan MJ (2018) Transient expression of CRISPR/Cas9 machinery targeting *TcNPR3* enhances defense response in *Theobroma cacao*. Front. Plant. Sci 9:329023. 10.3389/fpls.2018.00268

21. Foley RC, Kidd BN, Hane JK, Anderson JP, Singh KB (2016) Reactive oxygen species play a role in the infection of the necrotrophic fungi, *Rhizoctonia solani*, in wheat. PLoS One 11(3):e0152548. 10.1371/journal.pone.0152548

22. Gao Y, Xue CY, Liu JM, He Y, Mei Q, Wei S, Xuan YH (2021) Sheath blight resistance in rice is negatively regulated by WRKY53 via *SWEET2a* activation. Biochem. Biophys. Res. Commun. 585:117–123. 10.1016/j.bbrc.2021.11.042

23. Hiei Y, Komari T (2008) *Agrobacterium*-mediated transformation of rice using immature embryos or calli induced from mature seed. Nature protoc. 3(5):824–834. 10.1038/nprot.2008.46

24. Jesudoss D, Ponnurangan V, Kumar MPR, Kumar KK, Mannu J, Sankarasubramanian H, Duraialagaraja S, Eswaran K, Loganathan A, Shanmugam V (2024) Advances in breeding, biotechnology, and nanotechnological approaches to combat sheath blight disease in rice. Mol. Biol. Rep. 51(1):958. 10.1007/s11033-024-09889-5

25. Jiang G, Yin D, Shi Y, Zhou Z, Li C, Liu P, Jia Y, Wang Y, Liu Z, Yu M, Wu X (2020) *OsNPR3.3*-dependent salicylic acid signaling is involved in recessive gene *xa5*-mediated immunity to rice bacterial blight. Sci. Rep. 10(1):6313. 10.1038/s41598-020-63059-8

26. Jun JH, Ha CM, Fletcher JC (2010) BLADE-ON-PETIOLE1 coordinates organ determinacy and axial polarity in *Arabidopsis* by directly activating ASYMMETRIC LEAVES2. Plant Cell 22(1):62–76. 10.1105/tpc.109.070763

27. Kauffman HE, Reddy APK, Hsieh SPY, Merca SD (1973) An improved technique for evaluating resistance of rice varieties to *Xanthomonas oryzae*. Plant Dis. Rep. 57:537–541.

28. Ke Y, Hui S, Yuan M (2017) *Xanthomonas oryzae* pv. *oryzae* inoculation and growth rate on rice by leaf clipping method. Bio. Protoc. 7(19):e2568. 10.21769/BioProtoc.2568

29. Kinkema M, Fan W, Dong X (2000) Nuclear localization of NPR1 is required for activation of PR gene expression. Plant cell 12(12):2339–2350. 10.1105/tpc.12.12.2339

30. Kouzai Y, Kimura M, Watanabe M, Kusunoki K, Osaka D, Suzuki T, Matsui H, Yamamoto M, Ichinose Y, Toyoda K, Matsuura T (2018) Salicylic acid-dependent immunity contributes to resistance against Rhizoctonia solani, a necrotrophic fungal agent of sheath blight, in rice and *Brachypodium distachyon*. New Phytol. 217(2):771–783. 10.1111/nph.14849

31. Kouzai Y, Kimura M, Yamanaka Y, Watanabe M, Matsui H, Yamamoto M, Ichinose Y, Toyoda K, Onda Y, Mochida K, Noutoshi Y (2016) Expression profiling of marker genes responsive to the defence-associated phytohormones salicylic acid, jasmonic acid and ethylene in *Brachypodium distachyon*. BMC plant biol. 16:1–11. 10.1186/s12870-016-0749-9

32. Lefevere H, Bauters L, Gheysen G (2020) Salicylic acid biosynthesis in plants. Front. Plant. Sci. 11:338. 10.3389/fpls.2020.00338

33. Li J, Mahajan A, Tsai MD (2006) Ankyrin repeat: a unique motif mediating protein−protein interactions. Biochem. 45(51):15168–15178. https://pubs.acs.org/doi/10.1021/bi062188q

34. Lin QJ, Chu J, Kumar V, Yuan DP, Li ZM, Mei Q, Xuan YH (2021) Protein phosphatase 2A catalytic subunit PP2A-1 enhances rice resistance to sheath blight disease. Front. Genome. Ed. 3:632136. 10.3389/fgeed.2021.632136

35. Liu H, Ding Y, Zhou Y, Jin W, Xie K, Chen LL (2017) CRISPR-P 2.0: an improved CRISPR-Cas9 tool for genome editing in plants. Mol. plant 10(3):530–532. 10.1016/j.molp.2017.01.003

36. Liu W, Xie X, Ma X, Li J, Chen J, Liu YG (2015) DSDecode: a web-based tool for decoding of sequencing chromatograms for genotyping of targeted mutations. Mol. plant 8(9):1431–1433. 10.1016/j.molp.2015.05.009

37. Livak KJ, Schmittgen TD (2001) Analysis of relative gene expression data using real-time quantitative PCR and the 2−^ΔΔC^T method. Methods 25(4):402–408. 10.1006/meth.2001.1262

38. Maier F, Zwicker S, Hueckelhoven A, Meissner M, Funk J, Pfitzner AJ, Pfitzner UM (2011) NONEXPRESSOR OF PATHOGENESIS-RELATED PROTEINS1 (NPR1) and some NPR1-related proteins are sensitive to salicylic acid. Mol. Plant Pathol. 12(1):73–91. 10.1111/j.1364-3703.2010.00653.x

39. Margani R, Widadi S (2018) Utilizing *Bacillus* to inhibit the growth and infection by sheath blight pathogen, Rhizoctonia solani in rice. IOP Conf. Ser. Earth Environ. Sci. 142:012070. 10.1088/1755-1315/142/1/012070

40. Mishra S, Roychowdhury R, Ray S, Hada A, Kumar A, Sarker U, Aftab T, Das R (2024) Salicylic acid (SA)-mediated plant immunity against biotic stresses: An insight on molecular components and signaling mechanism. Plant Stress :100427. 10.1016/j.stress.2024.100427

41. Molla KA, Karmakar S, Chanda PK, Ghosh S, Sarkar SN, Datta SK, Datta K (2013) Rice oxalate oxidase gene driven by green tissue-specific promoter increases tolerance to sheath blight pathogen (*Rhizoctonia solani*) in transgenic rice. Mol Plant Pathol. 14(9):910–922. 10.1111/mpp.12055

42. Molla KA, Karmakar S, Chanda PK, Sarkar SN, Datta SK, Datta K (2016) Tissue-specific expression of *Arabidopsis* NPR1 gene in rice for sheath blight resistance without compromising phenotypic cost. Plant Sci. 250:105–114. 10.1016/j.plantsci.2016.06.005

43. Molla KA, Karmakar S, Molla J, Bajaj P, Varshney RK, Datta SK, Datta K (2020) Understanding sheath blight resistance in rice: the road behind and the road ahead. Plant Biotechnol. J. 18(4):895–915. 10.1111/pbi.13312

44. Moon SJ, Park HJ, Kim TH, Kang JW, Lee JY, Cho JH, Lee JH, Park DS, Byun MO, Kim BG, Shin D (2018) *OsTGA2* confers disease resistance to rice against leaf blight by regulating expression levels of disease related genes via interaction with NH1. PLoS One 13(11):e0206910. 10.1371/journal.pone.0206910

45. Naeimi S, Khosravi V, Varga A, Vágvölgyi C, Kredics L (2020) Screening of organic substrates for solid-state fermentation, viability and bioefficacy of *Trichoderma harzianum* AS12-2, a biocontrol strain against rice sheath blight disease. Agron. 10(9):1258. 10.3390/agronomy10091258

46. Park DS, Sayler RJ, Hong YG, Nam MH, Yang Y (2008) A method for inoculation and evaluation of rice sheath blight disease. Plant Dis. 92(1):25–29. 10.1094/PDIS-92-1-0025

47. Peng X, Wang H, Jang JC, Xiao T, He H, Jiang D, Tang X (2016) *OsWRKY80*-*OsWRKY4* module as a positive regulatory circuit in rice resistance against *Rhizoctonia solani*. Rice 9:1–14. 10.1186/s12284-016-0137-y

48. Ponnurangan V, Namachivayam R, Pradeep RK, Jesudoss D, Eswaran K, Loganathan A, Kumar KK, Vaikuntavasan P, Maduraimuthu D, Shanmugam V (2025) Biotechnological breakthroughs in rice disease management: an overview from transgenics to CRISPR. Mol. Biol. Rep. 52(1):616. 10.1007/s11033-025-10701-1

49. Porebski S, Bailey LG, Baum BR (1997) Modification of a CTAB DNA extraction protocol for plants containing high polysaccharide and polyphenol components. Plant Mol. Biol. Rep. 15:8–15. 10.1007/BF02772108

50. Rochon A, Boyle P, Wignes T, Fobert PR, Després C (2006) The coactivator function of *Arabidopsis* NPR1 requires the core of its BTB/POZ domain and the oxidation of C-terminal cysteines. Plant Cell 18(12):3670–3685. 10.1105/tpc.106.046953

51. Senapati M, Tiwari A, Sharma N, Chandra P, Bashyal BM, Ellur RK, Bhowmick PK, Bollinedi H, Vinod KK, Singh AK, Krishnan SG (2022) *Rhizoctonia solani* Kühn pathophysiology: Status and prospects of sheath blight disease management in rice. Front. Plant Sci. 13:881116. 10.3389/fpls.2022.881116

52. Shah J (2003) The salicylic acid loop in plant defense. Curr. Opin. Plant. Biol. 6(4):365–371. 10.1016/S1369-5266(03)00058-X

53. Shanmugam V, Wang YW, Tsednee M, Karunakaran K, Yeh KC (2015) Glutathione plays an essential role in nitric oxide-mediated iron-deficiency signaling and iron-deficiency tolerance in *Arabidopsis*. Plant J. 84(3):464–477. 10.1111/tpj.13011

54. Shanthinie A, Vignesh P, Kumar KK, Arul L, Varanavasiappan S, Manonmani S, Jeyakumar P, Kokiladevi E, Sudhakar D (2024) Enhancing rice grain quality through the knock-out of the *OsSPL16* gene. Plant Physiol. Rep. 29(2):308–315. 10.1007/s40502-024-00790-8

55. Son S, Moon SJ, Kim H, Lee KS, Park SR (2021) Identification of a novel NPR1 homolog gene, *OsNH5N16*, which contributes to broad-spectrum resistance in rice. Biochem. Biophys. Res. Commun. 549:200–206. 10.1016/j.bbrc.2021.02.108

56. Spoel SH, Koornneef A, Claessens SM, Korzelius JP, Van Pelt JA, Mueller MJ, Buchala AJ, Métraux JP, Brown R, Kazan K, Van Loon LC (2003) NPR1 modulates cross-talk between salicylate-and jasmonate-dependent defense pathways through a novel function in the cytosol. Plant Cell 15(3):760–770. 10.1105/tpc.009159

57. Sree DB, Shanthinie A, Vignesh P, Varanavasiappan S, Kumar KK, Arul L, Kokiladevi E, Manonmani S, Saranya N, Sudhakar D (2023) Targeted editing of *OsSWEET11* promoter for imparting bacterial leaf blight resistance in rice. Electron. J. Plant Breed. 14(3):938–947. 10.37992/2023.1403.106

58. Tiwari M, Srivastava S, Singh PC, Mishra AK, Chakrabarty D (2020) Functional characterization of tau class glutathione-S-transferase in rice to provide tolerance against sheath blight disease. 3Biotech 10:1-7. 10.1007/s13205-020-2071-3

59. Torres MA, Jones JD, Dangl JL (2006) Reactive oxygen species signaling in response to pathogens. Plant Physiol. 141(2):373–378. 10.1104/pp.106.079467

60. Wang L, Liu H, Yin Z, Li Y, Lu C, Wang Q, Ding X (2022) A novel guanine elicitor stimulates immunity in Arabidopsis and rice by ethylene and jasmonic acid signaling pathways. Front. Plant Sci. 13:841228. 10.3389/fpls.2022.841228

61. Willocquet L, Elazegui FA, Castilla N, Fernandez L, Fischer KS, Peng S, Teng PS, Srivastava RK, Singh HM, Zhu D, Savary S (2004) Research priorities for rice pest management in tropical Asia: a simulation analysis of yield losses and management efficiencies. Phytopathology 94(7):672–682. 10.1094/PHYTO.2004.94.7.672

62. Xie K, Minkenberg B, Yang Y (2015) Boosting CRISPR/Cas9 multiplex editing capability with the endogenous tRNA-processing system. Proc. Natl. Acad. Sci. 112(11):3570–3575.10.1073/pnas.1420294112

63. Xie X, Ma X, Zhu Q, Zeng D, Li G, Liu YG (2017) CRISPR-GE: a convenient software toolkit for CRISPR-based genome editing. Mol. plant 10(9):1246–1249. 10.1016/j.molp.2017.06.004

64. Xu RF, Li H, Qin RY, Li J, Qiu CH, Yang YC, Ma H, Li L, Wei PC, Yang JB (2015) Generation of inheritable and “transgene clean” targeted genome-modified rice in later generations using the CRISPR/Cas9 system. Sci. Rep. 5(1):11491. 10.1038/srep11491

65. Yang Y, Qi M, Mei C (2004) Endogenous salicylic acid protects rice plants from oxidative damage caused by aging as well as biotic and abiotic stress. Plant J. 40(6):909–919. 10.1111/j.1365-313X.2004.02267.x

66. Yu D, Chen C, Chen Z (2001) Evidence for an important role of WRKY DNA binding proteins in the regulation of NPR1 gene expression. Plant Cell 13(7):1527–1540. 10.1105/TPC.010115

67. Yuan Y, Zhong S, Li Q, Zhu Z, Lou Y, Wang L, Wang J, Wang M, Li Q, Yang D, He Z (2007) Functional analysis of rice NPR1-like genes reveals that *OsNPR1/NH1* is the rice orthologue conferring disease resistance with enhanced herbivore susceptibility. Plant Biotechnol. J. 5(2):313–324. 10.1111/j.1467-7652.2007.00243.x

68. Zhang Y, Cheng YT, Qu N, Zhao Q, Bi D, Li X (2006) Negative regulation of defense responses in *Arabidopsis* by two NPR1 paralogs. Plant J. 48(5):647–656. 10.1111/j.1365-313X.2006.02903.x

69. Zhang Y, Fan W, Kinkema M, Li X, Dong X (1999) Interaction of NPR1 with basic leucine zipper protein transcription factors that bind sequences required for salicylic acid induction of the PR-1 gene. Proc. Natl. Acad. Sci. USA 96(11):6523–6528. 10.1073/pnas.96.11.6523

70. Zhang Z, Chen Y, Li B, Chen T, Tian S (2020) Reactive oxygen species: A generalist in regulating development and pathogenicity of phytopathogenic fungi. Comput. Struct. Biotechnol. J. 18:3344–3349. 10.1016/j.csbj.2020.10.024

71. Zheng A, Lin R, Zhang D, Qin P, Xu L, Ai P, Ding L, Wang Y, Chen Y, Liu Y, Sun Z (2013) The evolution and pathogenic mechanisms of the rice sheath blight pathogen. Nat. commun. 4(1):1424. 10.1038/ncomms2427

72. Zhou JM, Trifa Y, Silva H, Pontier D, Lam E, Shah J, Klessig DF (2000) NPR1 differentially interacts with members of the TGA/OBF family of transcription factors that bind an element of the PR-1 gene required for induction by salicylic acid. Mol. Plant Microbe Interact. 13(2):191–202. 10.1094/MPMI.2000.13.2.191

73. Zhou Y, Xu S, Jiang N, Zhao X, Bai Z, Liu J, Yao W, Tang Q, Xiao G, Lv C, Wang K (2022) Engineering of rice varieties with enhanced resistances to both blast and bacterial blight diseases via CRISPR/Cas9. Plant Biotechnol. J. 20(5):876–885. 10.1111/pbi.13766

